# Loss of endogenous tau suppresses APOE4-induced patterned behavioral decline and axon dysmorphia in a *C. elegans* model of Alzheimer’s disease

**DOI:** 10.1101/2025.05.06.652574

**Authors:** Eric A. Cardona, Chelsea J. Webber, Zheng Wu, Blythe M. Bolton, Elif Sarinay-Cenik, Jonathan T. Pierce

**Affiliations:** Department of Neuroscience, Center for Learning and Memory, University of Texas at Austin, Austin, TX; Department of Molecular Biosciences, University of Texas at Austin, Austin, TX

## Abstract

Alzheimer’s disease (AD) causes a characteristic spatiotemporal pattern of neurodegeneration. The factors that account for this pattern of degeneration are unclear. Previously, we generated a model of AD using the nematode *Caenorhabditis elegans* with the AD risk variant of apolipoprotein E, *APOE4*. We showed that HSN class neurons degenerate in early adult animals. Here, we perform behavioral analyses to deduce the effect of APOE4 on the function of distinct neuronal circuits. We found evidence that APOE4 induces dysfunction of other neurons; this spatiotemporal pattern roughly correlates with endogenous levels of PTL-1, the *C. elegans* homolog of human MAPT also known as tau. Moreover, deletion of *ptl-1* suppressed defects in multiple behaviors, suggesting broad protective effects across the nervous system including the HSN neurons. Lastly, we show that PTL-1 in the touch receptor neurons, where PTL-1 is most abundant, contributes non-cell autonomously for age-related axon dysmorphia and dysfunction of the HSN neurons. Our results suggest that *C. elegans* may provide a useful *in vivo* system to study how endogenous tau acts downstream of APOE4 to cause progressive, patterned neurodegeneration.

## Introduction

Neurodegenerative disorders such as Alzheimer’s disease (AD), Parkinson’s disease (PD), and frontotemporal lobar dementia (FTLD) present with characteristic spatiotemporal patterns of neuronal cell death (Braak and Braak, 1991; Bahia et al., 2013; Reviewed in Rietdijk et al., 2017). Degeneration in characteristic brain regions corresponds to loss of brain region-associated faculties, such as memory in AD (Rosen et al., 2005). Genetic risk factors have been previously associated with AD. Among them, the ε4 variant of Apolipoprotein E (APOE4) is the most common genetic risk factor for earlier onset and faster progression of sporadic AD and AD-like pathology in individuals with Down syndrome (Larsen et al. 2024; Serrano-Pozo et al., 2021). A recent study proposed that *APOE4* homozygosity itself may be a single-gene driver of AD, akin to mutation of the familial AD genes *APP*, *PSEN1*, and *PSEN2* (Fortea et al., 2024; Ryan and Rossor, 2010). Moreover, APOE4 has been shown to increase the severity of other neurodegenerative diseases such as PD and FTLD (Robinson et al., 2018; Koriath et al., 2019; Davis et al., 2020). How APOE4 worsens neurodegeneration (as in AD, PD, and FTLD) or, more interestingly, may induce it (as suspected in AD) and contribute to patterned degeneration is unclear.

As *APOE4* homozygosity alone may cause AD, it is critical to understand how APOE4 contributes to patterned neurodegeneration. One hypothesis is APOE4-induced neurodegeneration follows a spatiotemporal pattern determined by cellular levels of specific vulnerability factors. Multiple lines of evidence suggest that the microtubule associated protein gene, *MAPT*, which encodes tau, represents one such vulnerability factor. First, pathogenic tau forms intracellular aggregates, and their accumulation is closely linked to neuronal dysfunction in AD and other degenerative disorders termed tauopathies (Trojanowski et al., 1993; Lin et al., 2019). Second, tau-related pathology is highly correlated with neurodegeneration according to Braak Staging, a well-established metric of how AD progresses with age (Braak and Braak, 1991; Köpke et al., 1993; Ghoshal et al., 2002). Third, transgenic expression of pathological tau is sufficient to cause neurodegeneration (Mocanu et al., 2008; Spittaels et al., 1999; www.alzforum.org/research-models). Fourth, depletion of endogenous tau mitigates cellular correlates of AD pathology in rodent models of AD that overexpress human *APP* (Roberson et al., 2007; Vossel et al., 2010) or human tau with aggregate-inducing mutations (Wegmann et al., 2015; DeVos et al., 2017). Fifth, growing evidence suggests that pathogenic tau protein can spread in a trans-synaptic fashion between brain regions *in vivo* with age (Liu et al., 2012; de Calignon et al., 2012; Iba et al., 2013; Fu et al., 2016; Meisl et al., 2021; Cornblath et al., 2021). Altogether, these and many other studies provide strong evidence for tau as a candidate vulnerability factor underlying patterned neurodegeneration. Progress has been made to uncover the relationship between APOE4, tau, and neurodegeneration (Shi et al., 2017; Therriault et al., 2020; Seo et al., 2023; Koutsodendris et al., 2023), but whether APOE4 is a causative factor in the pathogenic spread of tau or how APOE4 may contribute to a combination of cell autonomous and non-cell autonomous processes is less clear. Additionally, because rodent models of AD often do not fully recapitulate the neurodegeneration observed in human patients, the cellular and molecular bases for this critical pathological phenotype have yet to be uncovered.

The nematode *Caenorhabditis elegans* has been famously leveraged to discover conserved *in vivo* mechanisms of cell death (Horvitz, 2003). It also represents a potentially useful system to study the cellular and molecular bases for patterned neurodegeneration *in vivo*. To model neurodegeneration, past studies have overexpressed human disease genes and investigated the health and function of specific neurons *in vivo* (Faber et al., 1999; Baskoylu et al., 2018; van Ham et al., 2008; Vaccaro et al., 2012b). In transgenic *C. elegans* models, progress has been made to investigate genes involved with AD, notably *MAPT* and *APP* (Kraemer et al., 2003; Treusch et al., 2011; Fatouros et al., 2012; Benbow et al., 2020; Sae-Lee et al., 2020). A powerful feature of *C. elegans* as a model organism is that each one of its 302 neurons is identifiable. However, previous studies did not take full advantage of this to investigate whether specific neurons differ in vulnerability due to tau. Thus, the potential of *C. elegans* to test spatiotemporal patterns of neurodegeneration remains underutilized.

Here, we found that APOE4 induces a spatiotemporal pattern of behavioral decline in *C. elegans* as inferred through the assessment of behaviors that reflect the activity of known neurons. This pattern roughly correlates with neuronal expression levels of *ptl-1*, the sole worm ortholog of *tau*. Complementing our indirect approach, we observed directly via live microscopy that APOE4 induces progressive, age-related HSN axon dysmorphia coincident with dysfunction of that neuron. Deletion of *ptl-1* suppressed both APOE4-induced behavioral defects and HSN axon dysmorphia. Genetic ablation of the six neurons most enriched with PTL-1—the touch receptor neurons (TRNs)—similarly suppressed HSN dysfunction; this was phenocopied by targeted depletion of PTL-1 in the TRNs. These results suggest a non-cell autonomous role for PTL-1 in APOE4-induced degeneration. Overall, our findings position the APOE4 worm model as a useful system to study how endogenous tau/PTL-1 participates in the cellular and molecular mechanisms underlying patterned neuronal dysmorphia or degeneration. This may aid in the development of new therapeutic measures against tauopathies, such as AD.

## Results

### APOE4 induces progressive, widespread behavioral defects in *C. elegans*

Our previous *C. elegans* study showed that pan-neuronal expression of APOE4 caused degeneration and dysfunction in a subset of neurons—the HSN pair—in early adulthood, while many others, such as VA and VB motor neurons, appeared healthy and functional (Sae-Lee et al., 2020). In that work, we classified HSN neurons as degenerated when their somas appeared shrunken or absent using a fluorescent reporter. Here, we extend this study by testing whether APOE4 induces a broader pattern of neuronal dysfunction, neurodegeneration, or both. To do this, we assayed multiple behaviors to probe the function of distinct neuronal subsets and circuits, reasoning that behavioral performance would reveal the health of the underlying neurons. We selected seven different behaviors that rely primarily, but not exclusively, on mostly distinct subsets of identified neuron classes (Fig. S1). Five behaviors we tested—*egg laying*, *pharyngeal pumping*, *gentle* and *harsh touch response*, and *locomotor gait transition*—have specific neurons that are critical for behavioral execution. These include: the HSN class neurons for egg laying behavior (Schafer, 2005); the MC and M3 neurons for pumping of the pharyngeal feeding organ (Avery and You, 2012); the six TRNs for gentle touch response (Chalfie and Sulston, 1981; Chalfie et al., 1985); the FLP and PVD neurons, with lesser contribution from redundant mechanoreceptive neurons and interneurons, for harsh touch response (Li et al., 2011); and lastly, the ADE and PDE dopaminergic neurons for the transition between swim and crawl locomotor gaits (Vidal-Gadea et al., 2011). We also assessed two readouts of locomotor behavior that rely on many more neurons with partial redundancy. *Short-term locomotion* relies on over 100 motor neurons and about one dozen pre-motor interneurons; *long-term locomotion* relies on those neurons and additional sensory neurons and interneurons to mediate switches between roaming and dwelling states (Flavell, et al., 2020).

We evaluated behavioral performance at adult days 1 through 5 (D1-5) (Fig. 1A left). This age range encompasses the peak reproductive period when *C. elegans* is active across a range of behaviors and lays eggs (Scharf et al., 2021). We refer to “age of onset” of behavioral decline as the first day of the adult stage that APOE4 worms displayed significantly impaired behavior compared to age-matched controls.

**Fig. 1.**
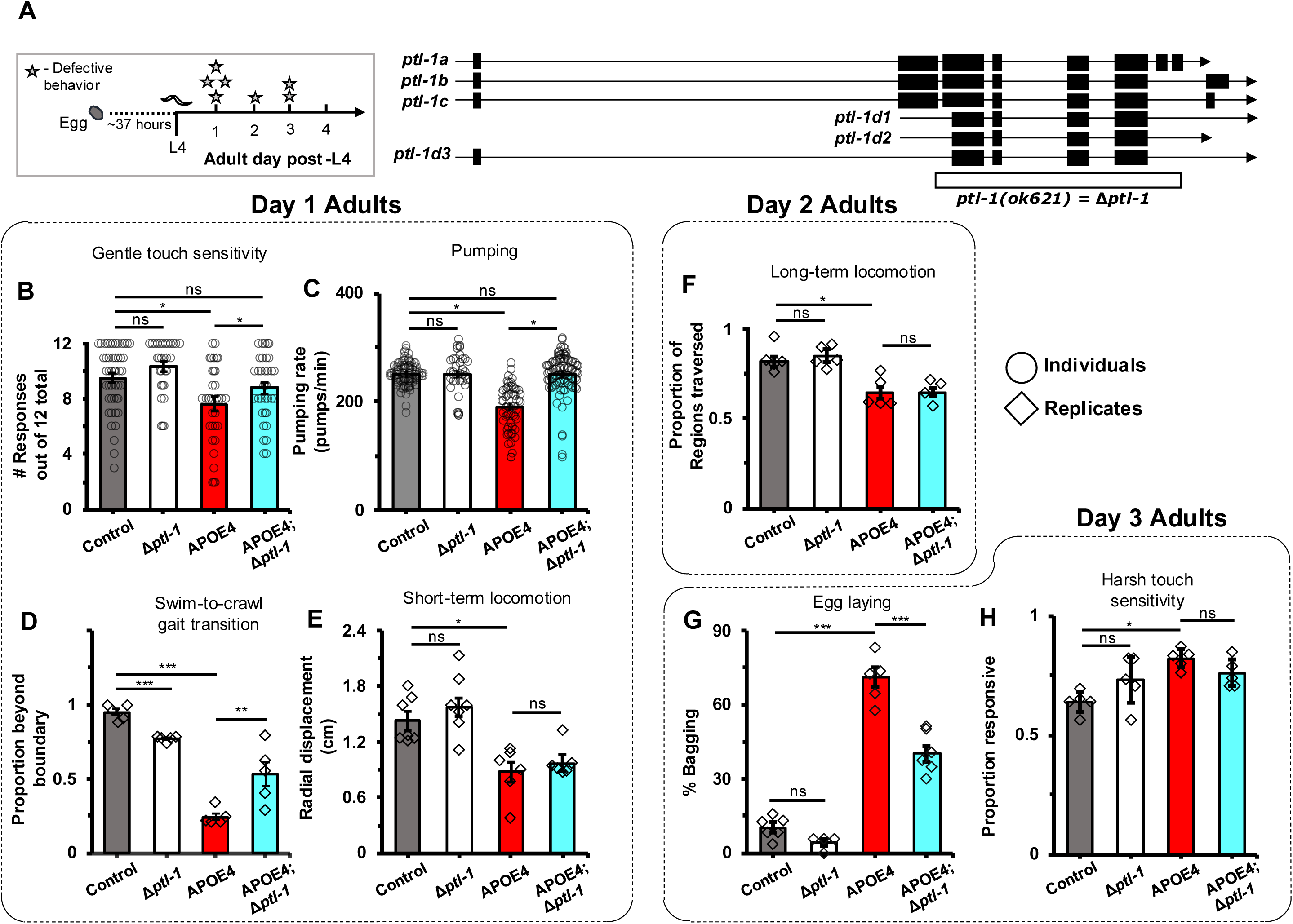
Deletion of *ptl-1* partially suppresses progressive pattern of behavioral decline induced by APOE4. (A left) Timeline to the adult age of onset of defective or abnormal behaviors in APOE4 strain. (A right) Endogenous *ptl-1* locus with all known isoforms and the portion deleted in the *ok621* allele as indicated below. (B-H) Results of behavioral assays, used to proxy neuronal dysfunction, are arranged by the age of onset for APOE4-induced defects. Only in the cases of gentle touch sensation and pumping behaviors (B,C) are APOE4 defects suppressed such that there are no significant differences between control and APOE4;Δ*ptl-1* groups. (B-E) D1 adult onset. (B) Subtle defect in gentle touch sensitivity which is fully suppressed by *ptl-1* deletion. N=47 for control, N=30 for Δ*ptl-1*, N=33 for APOE4, and N=35 for APOE4;Δ*ptl-1*. (C) Subtle defect in pharyngeal pumping which is fully suppressed by *ptl-1* deletion. N=94 for control, N=37 for Δ*ptl-1*, N=61 for APOE4, and N=91 for APOE4;Δ*ptl-1*. (D) Severe defect in swim-to-crawl gait transition which is partially suppressed by *ptl-1* deletion. N=164 control, N=136 for Δ*ptl-1*, N=162 for APOE4, N=165 for APOE4;Δ*ptl-1*. (E) Subtle defect in short-term locomotion which is not suppressed by *ptl-1* deletion. N=47 for control, N=38 for Δ*ptl-1*, N=44 for APOE4, and N=44 for APOE4;Δ*ptl-1*. (F) D2 adult onset. Subtle defect in long-term locomotion was not suppressed by *ptl-1* deletion. N=31 control, N=32 for Δ*ptl-1*, N=28 for APOE4, and N=29 for APOE4;Δ*ptl-1*. (G,H) D3 adult onset. (G) Severe egg laying defect was partially suppressed by *ptl-1* deletion. N=196 for control, N=134 for Δ*ptl-1*, N=214 for APOE4, N=243 for APOE4;Δ*ptl-1*. (H) Subtle, abnormal increase in harsh touch sensitivity. N=25 for control, N=17 for Δ*ptl-1*, N=26 for APOE4, and N=31 for APOE4;Δ*ptl-1*. All data are represented as mean ± SEM. N is the total sample size representing all animals across replicates. Statistical comparisons were made using χ^2^ tests of independence for D,G and planned tests after two-way ANOVA for B,C,E,F,G. *: p<0.05; **: p<0.01; ***: p<0.001.

Over the course of D1-5 of adult, we found that APOE4 worms displayed differences in all seven behaviors compared to control (Fig. 1B-H). In as early as D1 adults, gentle touch sensitivity, pharyngeal pumping, and short-term locomotion performances decreased by 11%, 13%, and 24%, respectively; gait transition decreased by a marked 59% (Fig. 1B-E). In D2 adults, long-term locomotion showed a 12% decline. We also replicated our previous finding of an egg-laying defect that leads to larvae hatching internally, a salient phenotype called “bagging” (Sae-Lee et al., 2020; Fig. 1G). APOE4 worms showed a salient 60% incidence of bagging, peaking at D3. Whereas other behavioral performances declined, harsh touch sensitivity oddly increased in APOE4 worms relative to control beginning at D3 (Figs. 1H, S2). Altogether, these results indicate that APOE4 induces age-related abnormalities in multiple behaviors that primarily rely on distinct subsets of neurons. Thus, the APOE4 worm exhibits a wider spatiotemporal pattern of neuronal dysfunction, beyond the HSN neuron pair, across early worm adult age.

The APOE4 worm exhibited a few defects in behaviors as early as D1 adults. To test whether this represents a defect that originates during or after development we evaluated gentle touch and pumping behaviors in L4 stage worms (∼24 hours younger than D1 adults) out to D3 of adulthood for a new set of worms (Figs. 1A left, 2). A two-way ANOVA was used with planned pairwise comparisons. Gentle touch and pumping behavior did not statistically differ between control and APOE4 groups (compare L4 timepoints in Fig. 2A to 2B and 2C to 2D; see Fig. S3 with ages plotted together). Throughout adult stage, gentle touch remained unchanged as expected into D1 and D3 for control (Pan et al., 2011), whereas sensitivity decreased in APOE4 (p<0.02). In contrast, while pumping rate in control worms increased as expected from L4 to D1 (Hosono et al., 1980; Huang et al., 2004; p<0.05 Fig. 2C), the rate remained unchanged in APOE4 worms (Fig. 2D). Thus, there are likely to exist mixed or multiple mechanistic underpinnings of APOE4-induced behavioral decline, such as age-related decline (as with gentle touch) or developmental defects (as with pharyngeal pumping).

**Fig. 2.**
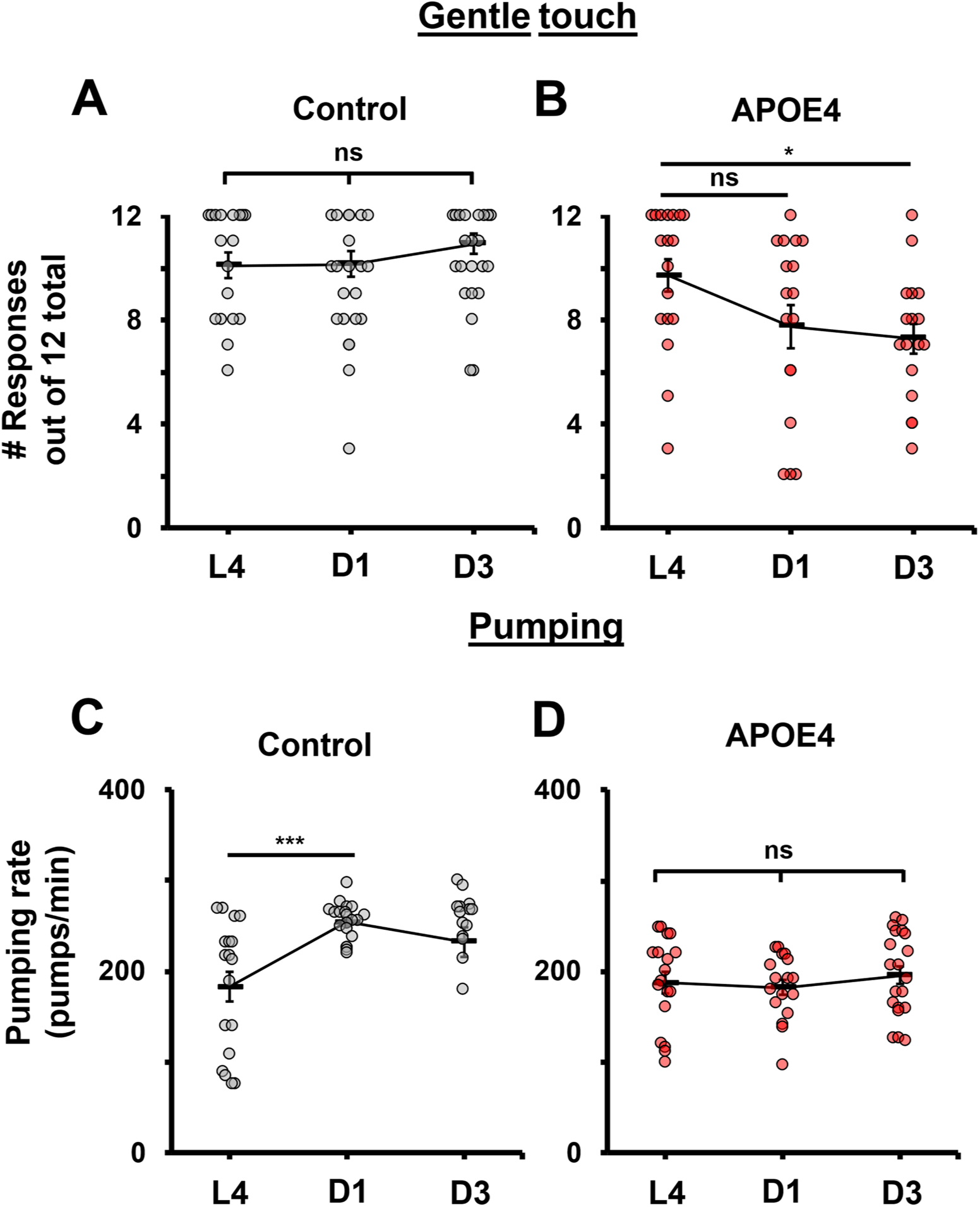
APOE4-induced behavioral decline may be the result of developmental and age-related defects. (A,B) Assays of gentle touch sensitivity. (A) Control worms maintained sensitivity from L4 (N=18) to D1 adult (N=22) to D3 adult (N=22) age. (B) APOE4 worms showed a decrease in sensitivity from L4 (N=18) to D1 adult (N=17) and remained unchanged afterward into D3 adult (N=17) age. (C,D) Assays of pharyngeal pumping. (C) Control worms showed an expected increase in pumping behavior from L4 (N=18) to D1 adult (N=20) that remained into D3 adult (N=16) age. (D) APOE4 worms showed no difference in pumping across L4 (N=18) to D1 adult (N=17) to D3 adult (N=20) ages. All data are represented as mean ± SEM. N is the total sample size. Statistical comparisons were made using planned tests after two-way ANOVA. *: p<0.05; **: p<0.01; ***: p<0.001.

### APOE4-induced progressive behavioral defects depend on the worm ortholog of tau

Above, we reported that gentle touch response is among the first behaviors to decline with APOE4 on D1 adult. This behavior is mediated by the six TRNs that feature a prominent array of microtubules that were exploited to discover the first microtubule gene (Chalfie and Thomson, 1979; Chalfie and Thomson, 1982). Studies have shown that tau binds to and stabilizes neuronal microtubules in mammals (Butner and Kirschner, 1991; Santarella et al., 2004). As one may expect from microtubule-enriched neurons, the worm homolog of tau, PTL-1, is highly enriched in the TRNs compared to other neurons, notably the PLM and ALM classes, according to recent single-cell transcriptomic analyses (Taylor et al., 2021; Roux et al., 2023; Fig. S4 green bars). We also noticed a rough correlation between endogenous levels of PTL-1 in specific neurons across the nervous system and the age of onset for APOE4-induced behavioral decline. This is depicted in Fig. S1, where gray shows what fraction of *ptl-1* is expressed in critical neurons that subserve each behavior. Given that tau represents a pathological hallmark associated with neurodegeneration in AD, we speculated that the high endogenous levels of PTL-1 in the TRNs may explain the early onset for the APOE4-induced gentle touch defect. Following this idea, we also hypothesized that deletion of *ptl-1* may broadly confer neuroprotection against APOE4. Into an APOE4 background we crossed a *ptl-1* mutant bearing the *ok621* deletion allele (Δ*ptl-1*). This allele represents a strong hypomorph, if not a null allele, as it lacks four exons that encode conserved microtubule-binding domains important for PTL-1 function (Goedert et al., 1996; Gordon et al., 2008; Hashi et al., 2015; Figs. 1A right, S5). Notably, the deleted region of PTL-1 contains residues conserved with mammalian tau that promote aggregation when phosphorylated (Köpke et al., 1993).

We found that deletion of *ptl-1* significantly suppressed APOE4-induced defects in four out of seven behaviors. Defects in gentle touch and pharyngeal pumping were completely suppressed (Fig. 1B,C), while defects in gait transition and egg laying were partly suppressed (Fig. 1D,G). We did not observe suppression of APOE4-induced abnormalities in short- and long-term locomotion nor for harsh touch sensitivity (Fig. 1E,F,H). As a control, we found that the *ptl-1* mutant was indistinguishable from control for six out of seven behaviors. Although the *ptl-1* mutant displayed a moderate defect in swim-to-crawl transition compared to control, the *ptl-1* mutant nonetheless partially suppressed the defect induced by APOE4 as opposed to additively worsening the defect (Fig. 1D). Also, although *ptl-1* mutant displayed a non-significant trend toward lower bagging compared to control animals, it significantly suppressed APOE4-induced bagging (Fig. 1G).

Altogether, we conclude that PTL-1 variably contributes to the neuronal dysfunction given multiple significant behavior changes induced by APOE4.

### APOE4-induced dysmorphia of HSN axon requires worm ortholog of tau

Previously, we showed that APOE4-induced bagging coincided with what had appeared to be degeneration of the bilaterally symmetric pair of HSN egg-laying motor neurons (Sae-Lee et al., 2020). Because deletion of *ptl-1* suppressed APOE4-induced bagging, we hypothesized that *ptl-1* deletion would also suppress HSN degeneration. To test this, we inspected the health of reporter-labeled HSN neurons in live worms with epifluorescent microscopy. Dorsal to the vulva, which is the egg-laying orifice, the HSNs are situated in the midbody of the worm (depicted in Fig. 3A). To quantify degeneration, we initially calculated various metrics of HSN soma geometry and reporter intensity across strains (Fig. S6A), focusing on D3 adult animals which is the age of onset for increased bagging (∼70% incidence in APOE4 worms; Figs. 1G, 3E).

**Fig. 3.**
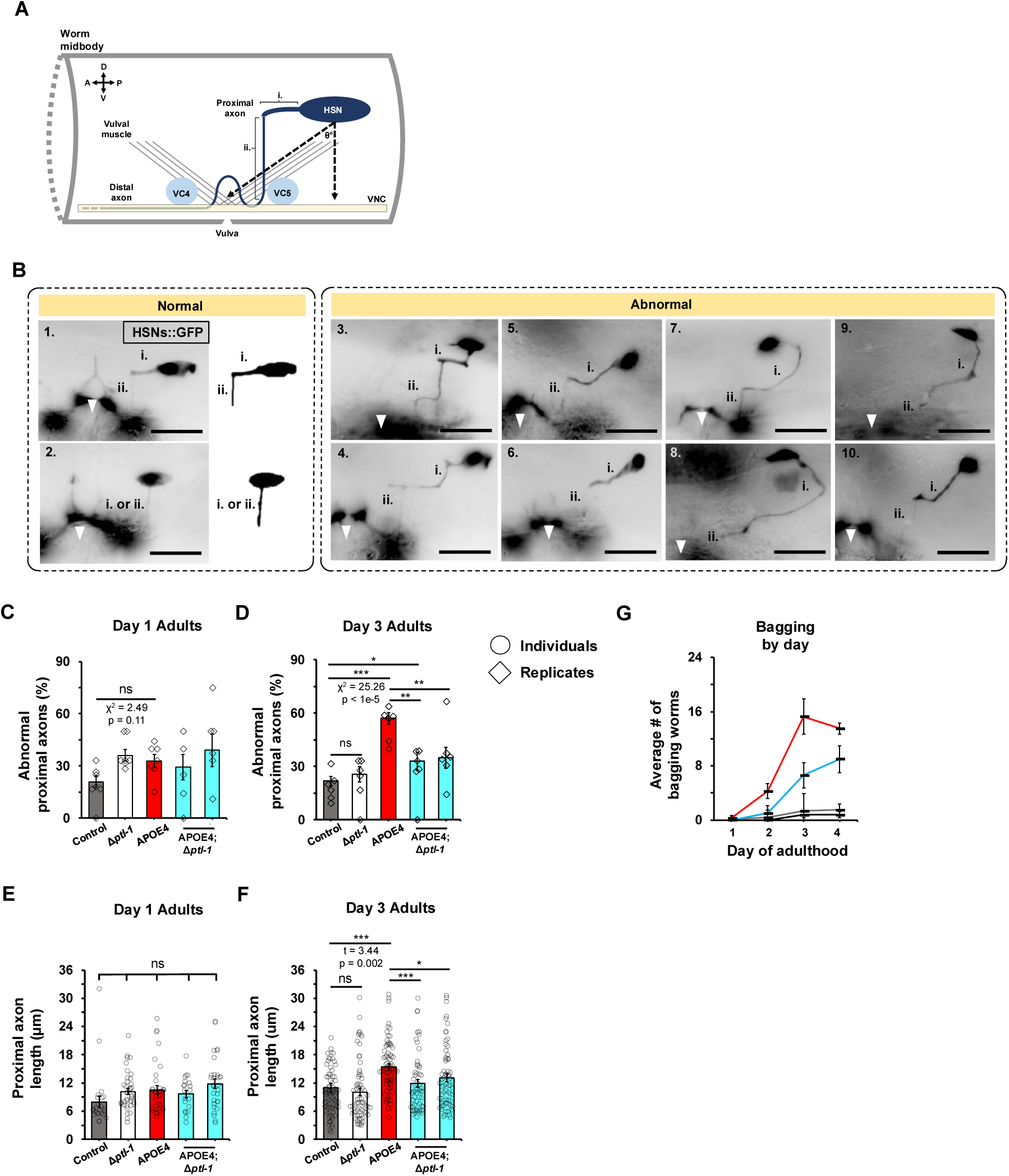
Deletion of *ptl-1* partially suppresses APOE4-induced HSN axon dysmorphia. (A) Cartoon of the worm midbody, centered about the vulva area, depicting: the HSN proximal axon, the VC4,5 neurons, vulval musculature, and ventral nerve cord (VNC). For the proximal axon: segment i. is the initial, anteriorly projecting portion and segment ii. is the ventrally projecting portion. (B) Representative live images of normal (1,2) and abnormal (3-10) HSN proximal axons visualized with GFP. Proximal axon segments are marked by i. and ii (with exception to image 2, where segment identity is less certain). The vulva area is also marked with a white arrowhead for anatomical reference. All scale bars are 20 μm. (C,D) Percentage of animals with abnormal HSN proximal axons in D1 and D3 adults. The baseline incidence of abnormal axons in control was similar between D1 and D3. (C) D1 adults. All other groups show non-significant, but slightly increased incidence of abnormal axons compared to control. N=37 for control, N=59 Δ*ptl-1*, N=51 for APOE4, N=22 for APOE4;Δ*ptl-1* (left), and N=37 for APOE4;Δ*ptl-1* (right). (D) The incidence of abnormal axons appears to correlate with defective egg laying behavior (See Fig. 1G) in D3 adults. N=101 for control, N=78 for Δ*ptl-1*, N=91 for APOE4, N=79 for APOE4;Δ*ptl-1* (left), and N=86 for APOE4;Δ*ptl-1* (right). (E,F) Quantification of HSN proximal axon lengths in D1 and D3 adults. The baseline axon length in control was similar between D1 and D3. (E) D1 adults. Axon length was similar across all groups. N=15 for control, N=13 Δ*ptl-1*, N=20 for APOE4, N=6 for APOE4;Δ*ptl-1* (left), and N=10 for APOE4;Δ*ptl-1* (right). (F) D3 adults. Axon length appears to correlate with the incidence of abnormal axons shown in (C) and defective egg laying behavior (See Fig. 1G). N=79 for control, N=66 for Δ*ptl-1*, N=72 for APOE4, N=54 for APOE4;Δ*ptl-1* (left), and N=66 for APOE4;*ptl-1* (right). (G) By-day breakdown of defective egg laying as quantified cumulatively in Fig. 1G. D3 data are replotted from Fig. 1G. All data are represented as mean ± SEM. N is the total sample size representing all animals across replicates. Statistical comparisons were made using χ^2^ tests of independence for C and D and planned comparisons after two-way ANOVAs for E and F. *: p<0.05; **: p<0.01; ***: p<0.001.

Consistent with our previous study (Sae-Lee et al., 2020), we found that the HSN soma appeared on average much dimmer in APOE4 than control worms, indicative of degeneration (Fig. S6D). However, deletion of *ptl-1* alone also appeared to have caused dimming along with abnormally large appearance of the HSN soma (Fig. S6B,D). We found no straightforward relationship between other soma-based morphometrics and HSN dysfunction as reflected by bagging incidence across control, *ptl-1*, and APOE4 strains (compare Fig. 1G to Fig. S6 measures). Thus, we conclude that HSN soma geometry and brightness are inconsistent predictors of HSN function in these strains, possibly due in part to unforeseen aspects of the fluorescent reporter (see Methods).

Seeking a more reliable cellular correlate of HSN function, we next investigated HSN axon morphology. Previous work using laser ablation and mutants showed the proximal axon portion (located between soma and vulval muscles)—but not distal portion which continues anteriorly—is required for egg laying behavior (Garriga et al., 1993; Fig. 3A). Therefore, we hypothesized that morphology of the proximal axon may serve as a useful correlate of HSN neuron function.

We visualized the HSN proximal axon with the same fluorescent reporter as above for the soma. The ∼11 μm-long HSN proximal axon typically comprises two segments joined at a right-angle (Fig. 3B image 1). The initial segment, which is qualitatively thicker, projects anteriorly from the soma (segment i.) before turning ventrally at a right-angle into a second, thinner segment (segment ii.) that joins the ventral nerve cord (VNC) and branches dorsally to innervate egg-laying musculature. The unbranched, distal axon continues anteriorly within the VNC to the head (Fig. 3A; Garriga et al., 1993). In our control reporter strain, we found that ∼58% of worms had a proximal axon with this typical two-segment, right-angle morphology (N=93; Fig. 3B image 1), while the remainder had a single ventrally-projecting proximal axon from the soma (Fig. 3B image 2). The latter single-segmented axons were thin and resembled segment ii. of two-segment HSN proximal axons. We speculate that the thicker, anteriorly directed segment i. did not develop in these worms because the HSN soma had already migrated to a position directly dorsal from the vulval muscles. Although the HSN proximal axon has been previously depicted with two segments connected at a right angle (Asakura et al., 2007; Tsutsui et al., 2021), both configurations have been described in the literature and are functional (Garriga et al., 1993). Thus, both types of proximal axons were scored as normal.

To check for axon abnormalities potentially caused by APOE4, we next qualitatively scored the morphology of HSN proximal axons and quantified their lengths in D3 adult animals. We found that the incidence of abnormal proximal axons was more than double in APOE4 compared to control worms. Proximal axons often took eccentric, elongated paths in APOE4 worms (compare Fig. 3B normal images 1,2 to abnormal images 3-10). This was not easily explained by abnormal positioning of the HSN soma, as the somas were similarly located in both APOE4 and control worms; soma positions were quantified as the HSN soma to VNC distance and the HSN soma to vulva area distance (Fig. S7C left) or the angle formed between these two vectors (Asakura et al., 2007; Fig. S7C right).

We observed at least three distinct qualitatively abnormal morphologies of the proximal axons relative to control. First, segment i. (Fig. 3B, image 3) or segment ii. (Fig. 3B, image 4) had an extra ∼90-degree turn. Second, segment i. connected to segment ii. with a typical right-angle turn, but displayed an atypical ∼45-degree portion, with an apparently wider segment near the soma (Fig. 3B images 5,6). Third, segments i. and ii. together took on a hook-like shape oriented anteriorly (Fig. 3B images 7,8). We also noted additional cases (<5% frequency) where these morphological abnormalities appeared to co-occur (e.g., segment ii. extra right-angle turn and segment i. 45-degree portion, Fig. 3B, image 10). Overall, we conclude that APOE4 appears to induce a variety of proximal axon abnormalities of the HSN neurons.

To help distinguish whether the HSN proximal axon abnormality represented a developmental defect, age-related degeneration, or both, we inspected younger adult animals to determine when the defect started. In D1 adults, there was no significant difference in the incidence of abnormal axons in APOE4 worms versus control (Fig. 3C). However, in D3 adults, the incidence of abnormal axons showed a three-fold increase in APOE4 worms versus control (Fig. 3D; see Fig. S8 with ages plotted together). Additionally, at the even younger L4 developmental stage, the entirety of HSN axon appears to project properly in ∼95% of APOE4 animals (Fig. S9). From these observations, we deduce that the proximal axon defect begins by at least D1 in some APOE4 worms and progresses in an age-related fashion, concurrent with the age of onset for increased bagging and with similar incidence (Fig. 3G). We thus conclude that the increased incidence in axon abnormality is likely to reflect a degenerative phenotype.

We next tested whether deletion of *ptl-1* might suppress APOE4-induced abnormal HSN proximal axon morphology. To partly control for genetic background, we inspected two independent isolates of APOE4;Δ*ptl-1* worms that were crossed with the HSN reporter strain. For both isolates, we found that the *ptl-1* deletion suppressed the APOE4-induced abnormal axon morphology (Fig. 3D).

The similar incidence of axon abnormality and bagging in APOE4 worms led us to conclude that abnormal proximal axons reflect a defect in HSN function (compare control to APOE4 in Figs. 1G, 3D,E). This is bolstered by our observation that deletion of *ptl-1* suppresses APOE4-induced defects in behavioral performance as well as HSN proximal axon morphology. Altogether, the above results support the idea that proximal HSN axon morphology is a reliably predictive correlate of efficient egg-laying behavior and thus HSN health and function.

Although HSN axons were abnormally shaped in our study, we did not observe consistent classical morphological signs of neuronal degeneration by D5 adulthood. Future studies should determine whether such degenerative features emerge at later ages.

### Ablating tau-rich touch receptor neurons protects HSN from APOE4-induced dysfunction

The spatiotemporal pattern of degeneration in AD is thought to be partly explained by pathological forms of tau spreading between connected neurons and across the brain (intra-and extra-synaptically; Liu et. al, 2012; de Calignon et al., 2012). As shown above, the systemic deletion of *ptl-1* protected against age-related APOE4-incuded behavioral defects and abnormal HSN proximal axon morphology. We next sought to test whether PTL-1 might contribute non-cell autonomously to degeneration, explaining in part the spatiotemporal pattern of behavioral decline in our APOE4 worm model. We hypothesized that targeting the neurons that are 1) most abundant with PTL-1, and 2) tend to become dysfunctional early, may suppress the subsequent degeneration of other neurons that express PTL-1 at a lower level. The TRNs represent the neuronal class most abundant in PTL-1 in *C. elegans*. Over 30% of neuronal *ptl-1* transcripts are expressed in the mere six TRNs (Taylor et al., 2021; Roux et al., 2023; Figs. S1,S4 green bars). Coincidentally, the TRNs also become dysfunctional in D1 adults (Fig. 1B). In contrast, the HSN neurons express ∼80% less *ptl-1* than the TRNs and become dysfunctional later starting in D3 adults (Figs. 1G,S3 blue bar; Taylor et al., 2021). Some TRNs are also nearby the HSN neurons, such as the PLMs which synapse onto the HSN neurons (White et al., 1986). Thus, the TRNs and HSN neurons in worm offer a convenient platform to test whether PTL-1 contributes non-cell autonomously to degeneration.

To assess the TRNs and PTL-1 protein *in vivo* during our targeted manipulations, we took advantage of a control strain in which endogenous PTL-1 protein is translationally tagged with mNeonGreen (PTL-1::mNG; Krieg, et al., 2017). Consistent with transcriptional studies (Taylor et al., 2021), TRNs were readily visible in living larval and adult stages by their abundant mNG-tagged PTL-1 (Representative PLM neuron in Figs. 4A2, 5B2). We also tested whether the mNG tag on PTL-1 impedes its apparent neurotoxic properties downstream of APOE4. If this were the case, we might expect less HSN dysfunction in APOE4;PTL-1::mNG worms. Instead, we found bagging incidence in APOE4;PTL-1::mNG worms (60%) was similar to that of the APOE4-alone strains generated in the current (∼70%) and our previous (50-60%) studies (note: all APOE4 strains depicted in Fig. 4A1 express mNG::PTL-1, versus those in Fig. 1G which do not; Sae-Lee et al., 2020). We conclude that the endogenous mNG tag does not interfere with the presumed toxicity of PTL-1 in an APOE4 background.

**Fig. 4.**
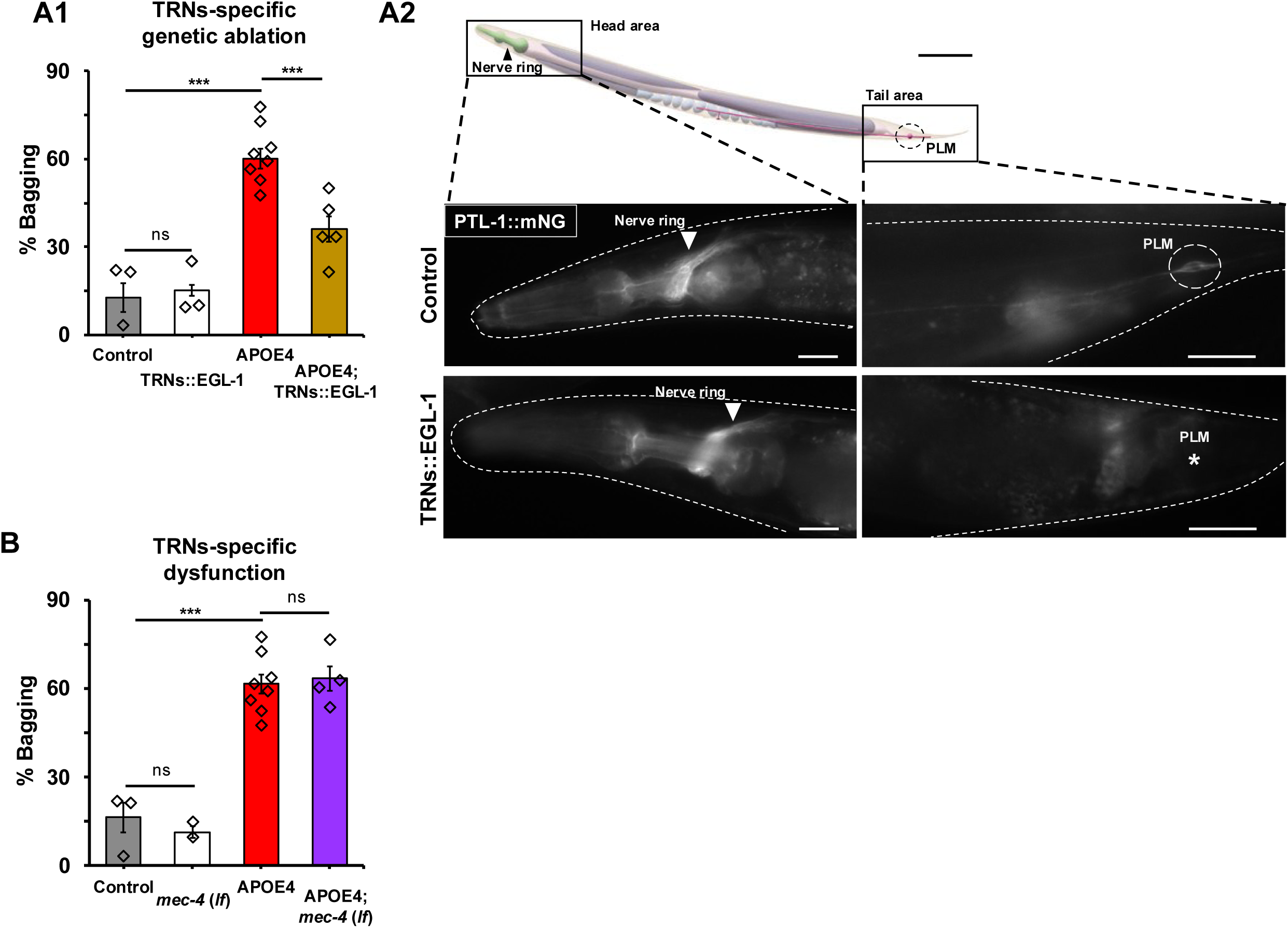
Targeted genetic ablation, but not abolished function, of the TRNs protects HSN against APOE4-induced dysfunction. (A1) Egg-laying behavior across groups. Genetic ablation of the TRNs partially suppressed egg-laying defects in APOE4 animals as with the *ptl-1* deletion (See Fig. 1G). N=104 for control, N=171 for TRNs::EGL-1, N=525 for APOE4, and N=190 for APOE4;TRNs::EGL-1. (A2 top) Cartoon depicting the head and tail areas wherein the nerve ring and PLM neuron, respectively, are located. (A2 bottom) Representative live images of nerve ring (arrowhead) and PLM neuron (within dashed circle if present; marked with asterisk if absent) visualized with PTL-1::mNG in control and TRNs::EGL-1 animals. Dashed borders indicate the worm body outline. All scale bars are 20 μm. (B) Egg-laying behavior across groups. In contrast to genetic ablation, abolishing TRN function did not suppress egg laying defects. N=104 for control, N=123 for *mec-4*(*lf*), N=525 for APOE4, and N=226 for APOE4;*mec-4*(*lf*). All data are represented as mean ± SEM. N is the total sample size representing all animals across replicates. All statistical comparisons were made using χ^2^ tests of independence. control and APOE4 data are re-plotted in A1 and B. *: p<0.05; **: p<0.01; ***: p<0.001.

For our first targeted manipulation, we tested whether ablation of the TRNs suppresses APOE4-induced HSN dysfunction. To do this, we leveraged the apoptotic trigger EGL-1, the sole worm homolog of the human BH3-only domain proteins BAM and BID. Targeted expression of *egl-1* has been previously used to kill specified cells (Chang et al., 2006). To kill the TRNs, we generated a strain whose *egl-1* expression is driven by the promoter of *mec-17*, a gene expressed exclusively in the TRNs (TRNs::EGL-1; Wu et al., 2025). The TRNs::EGL-1 strain was then crossed into the mNG-tagged *ptl-1* genetic background. To confirm the efficacy of TRN genetic ablation, we first assayed gentle touch responses; we found that only 10% were touch-responsive compared to control (N = 50). Second, we took advantage of the relative abundance of mNG-tagged PTL-1 in the TRNs that could be directly observed using epifluorescence microscopy (Fig. 4A2 control). Anatomically, this was easier to test with the PLM class of TRNs due to how its soma and process are confined in the narrow tail region, in contrast with the other TRN classes with soma situated in the thicker midbody (Fig. 4A2 control). We found that 93% of TRNs::EGL-1 animals had no detectable PTL-1::mNG where PLM soma and processes were expected compared to control (N=46; Fig. 4A2). Third, we confirmed ablation by imaging the PLM cell body with Differential Interference Contrast (DIC) optics (Fig. S10). Together, these results suggest a successful cell ablation in most cases.

We found that ablating the TRNs partially suppressed APOE4-induced HSN dysfunction as indicated by a ∼30% reduction in bagging incidence (Fig. 4A1 yellow bar). Ablation of the TRNs alone in the control strain had no effect on bagging (Fig. 4A1 white bar). Impressively, ablating the TRNs conferred a similar magnitude of protection as afforded by systemic deletion of *ptl-1* (compare blue bar in Fig. 1G to yellow bar in Fig. 4A1). Thus, we deduce that TRNs activity and/or their cellular contents may contribute non-cell autonomously to APOE4-induced HSN degeneration.

Because TRN activity is obviously lost after genetic ablation, we asked whether simply blocking TRNs function alone as opposed to whole cell ablation may protect the HSNs against APOE4. We tested this by using a deletion allele of *mec-4*, a gene that encodes mechanoreceptive DEG/ENaC channel expressed predominantly in the TRNs (Chalfie and Sulston, 1981). We took advantage of the *u253* loss-of-function allele of *mec-4*, referred hereon as *mec-4*(*lf*), abolishes transduction of mechanosensory currents and gentle touch response yet leaves the TRNs otherwise physiologically and morphologically normal (O’Hagan, et al., 2005). We found that *mec-4*(*lf*) failed to suppress APOE4-induced HSN dysfunction (Fig. 4B). From this, we conclude that functional impairment of the TRNs is insufficient to protect HSNs against APOE4.

### Targeted reduction of tau in touch receptor neurons protects HSN against APOE4

Because the worm ortholog of tau, PTL-1, is especially abundant in the TRNs (Fig. S4 green bars), we hypothesized that targeted depletion of PTL-1 contained in the TRNs would protect against APOE4-induced HSN dysfunction, phenocopying TRNs ablation. To test this, we knocked down *ptl-1* expression in the TRNs using cell-selective RNAi (TRNs::*ptl-*1 KD; Esposito et al., 2007). To determine the efficacy and selectivity of the knockdown, we quantified PTL-1::mNG fluorescence intensity in TRN soma versus example non-targeted neurons. Conveniently, the soma of the non-targeted VC4 and VC5 class motor neurons are also easily detectable by PTL-1::mNG fluorescence (Fig. 3A blue neurons; Fig. S4 blue bars). Compared to control, TRNs::*ptl-1* KD worms had an average 40% decrease in PTL-1::mNG fluorescence in five out of the six TRNs (Fig. 5A2). In contrast, PTL-1::mNG fluorescence intensity was unchanged in the non-targeted VC4 and VC5 neurons (Fig. 5A2). These observations suggested an effective and selective knockdown of *ptl-1* in the TRNs.

**Fig. 5.**
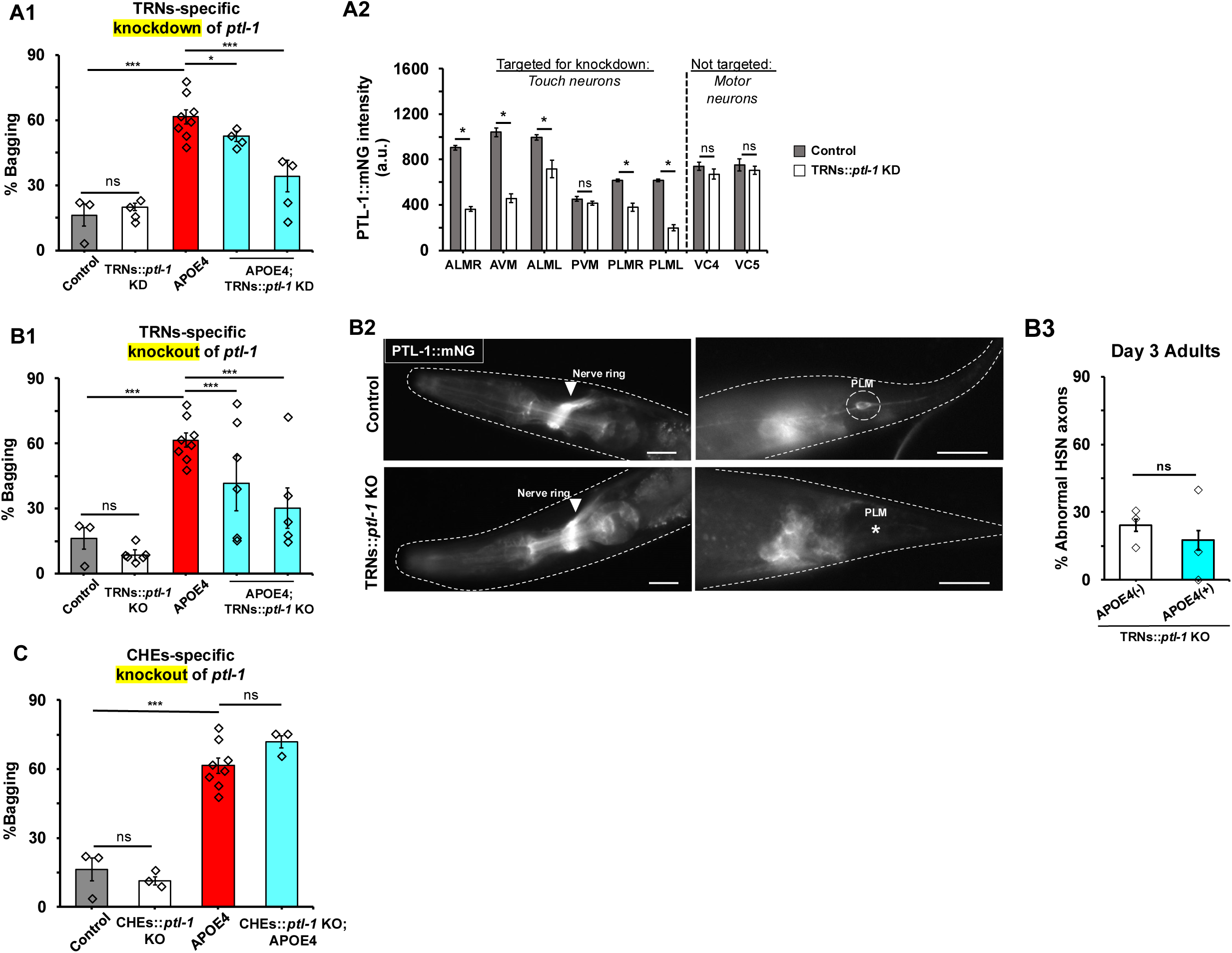
Targeted depletion of PTL-1 in the TRNs protects HSN against APOE4-induced dysfunction. (A1) Egg-laying behavior across groups. N=104 for control, N=180 for TRNs::*ptl-1* KD, N=525 for APOE4, N=171 for APOE4;TRNs::*ptl-1* KD (left), and N=108 for APOE4;TRNs::*ptl-1* KD (right). (A2) Quantification of *ptl-1* knockdown in the targeted TRNs and non-targeted VCs4,5 neurons between control and TRNs::*ptl-1* KD animals using PTL-1::mNG fluorescence intensity in the soma. (B1) Egg-laying behavior across groups. N=104 for control, N=348 for TRNs::*ptl-1* KO, N=525 for APOE4, N=423 for APOE4;TRNs:: *ptl-1* KO (left), and N=519 for APOE4;TRNs:: *ptl-1* KO (right). (B2) Representative live images of nerve ring (arrowhead) and PLM neuron (within dashed circle if present; area marked with asterisk if absent) visualized with PTL-1::mNG in control and TRNs::*ptl-1* KO animals. Dashed borders indicate the worm body outline. All scale bars are 20 μm. (B3) Percentage of animals with abnormal HSN proximal axons in D3 adults using independent reporter transgenics from Fig. 3. N=31 for TRNs::*ptl-1* KO and N=17 for APOE4;TRNs::*ptl-1* KO. (C) Egg-laying behavior across groups. N=104 for control, N=80 for CHEs::*ptl-1* KO, N=525 for APOE4, and N=78 for APOE4;TRNs:: *ptl-1* KO. All data are represented as mean ± SEM. N is the total sample size representing all animals across replicates. Statistical comparisons were made using χ^2^ tests of independence for A1,B1,B3,C and Student’s t-test for A2. Control and APOE4 data are re-plotted in A1, B1, and C. *: p<0.05; **: p<0.01; ***: p<0.001.

Using this strategy, we found that knockdown of *ptl-1* in the TRNs partially suppressed APOE4-induced HSN dysfunction. This was determined after scoring bagging in two independent isolates of APOE4 worms crossed with the TRNs::*ptl-1* KD strain to partly control for genetic background (Fig. 5A1 blue bars). In contrast, knockdown of *ptl-1* in TRNs alone in the control strain had no effect on bagging (Fig. 5A1 white bar). While significant, the suppressive effect was noticeably weaker compared to the systemic *ptl-1* deletion (compare with APOE4;Δ*ptl-1* strain, blue bar in Fig. 1G). We speculated that the weaker suppression may be explained by RNAi reducing only a portion of *ptl-1* transcripts in the TRNs (Fig. 5A2). This prompted us to test whether complete loss of *ptl-1* in the TRNs more strongly suppresses HSN dysfunction than in APOE4;TRNs::*ptl-1* KD worms.

To achieve a targeted *ptl-1* knockout in the TRNs, we used an intersectional Cre-LoxP approach. *LoxP* sites were first introduced via CRISPR to flank the endogenous *ptl-1*::*mNG* locus (*LoxP*::*ptl-1*::*LoxP* strain). *LoxP*::*ptl-1*::*LoxP* worms were then crossed with a strain that expresses Cre recombinase under the same TRNs-specific promoter as our TRNs::EGL-1 strain (Harterink et al., 2018; Sanfeliu-Cerdán et al., 2023; Wu et al., 2025). This should result in a selective knockout of *ptl-1* in the TRNs (TRNs::*ptl-1* KO). The efficacy of the *ptl-1* knockout was confirmed first by evaluating the presence of PTL-1 in the PLM neurons via PTL-1::mNG fluorescence signal (Fig. 5B2). PTL-1::mNG was absent in 100% of PLM neurons in TRNs::*ptl-1* KO animals (N=45). Second, we confirmed the loss of the PLM cell body itself using Normarskii optics (Fig. S10). Evaluation of protein by Western blot with the FLAG epitope on the tag using whole-animal lysate was inconclusive due to high variability (Krieg et al., 2017; Fig. S11).

As with the knockdown, we found that knockout of *ptl-1* in the TRNs partially suppressed APOE4-induced HSN dysfunction. This was determined after scoring bagging in two independent isolates of APOE4 worms crossed with the TRNs::*ptl-1* KO strain to partially control for genetic background (Fig. 5B1 blue bars). In contrast to the mild suppression after *ptl-1* knockdown (Fig. 5A1 blue bars), the average extent of suppression by knockout in the TRNs was more equivalent to that of the systemic deletion of *ptl-1* (Fig. 1G).

Finally, we inspected HSN axons directly to test whether knockout of *ptl-1* in the TRNs suppresses APOE4-induced abnormal morphology. To do this, we generated new transgenic reporter lines using our TRNs::*ptl-1* KO and APOE4;TRNs::*ptl-1* KO strains (Fig. 5B1 white and left blue bar). In these strains, we observed no difference in the incidence of abnormal HSN proximal axons comparing the no-APOE4 control to the APOE4 worms (Fig. 5B3). We conclude that knockout of *ptl-1* in the TRNs is sufficient to suppress the axon morphology defects induced by APOE4.

Does loss *ptl-1* in neurons other than the TRNs suppress APOE4-induced HSN dysfunction? Aside from the TRNs, other sensory neurons in *C. elegans* are also abundant in tau/PTL-1. These include a subset of over two dozen neurons that express the intraflagellar transport protein CHE-2 in the ciliary basal body (Fujiwara et al., 1999; Taylor et al., 2021; Fig. S4 yellow bars). As with our strategy for the TRNs, we generated a strain in which *ptl-1* was knocked out in the these sensory neurons by crossing our *LoxP*::*ptl-1*::*LoxP* strain a strain whose Cre recombinase expression is driven by the *<u>che</u>-2* promoter (CHEs::*ptl-1* KO). We observed no significant suppression in APOE4;CHEs::*ptl-1* KO compared to APOE4 alone (Fig. 5C), suggesting PTL-1 in the broad set of *che-2*-expressing sensory neurons is not required for APOE4-induced HSN dysfunction.

Altogether, we conclude that PTL-1 in the TRNs contributes non-cell autonomously to APOE4-induced dysmorphia of the HSN neuron axons. We do not conclude that TRNs represent the only source of harmful tau.

## Discussion

### APOE4-induced pattern of behavioral decline in *C. elegans*

Here, we build on our prior work by characterizing APOE4-induced, age-related abnormalities across seven behaviors in a *C. elegans* model of AD. The decline in behaviors provides evidence for a spatiotemporal APOE4-induced pattern of neuronal dysfunction and dysmorphia. Finally, we demonstrate that endogenous tau contributes to APOE4-induced axon dysmorphia and multiple behavioral defects.

*C. elegans* has served as a leading model organism to study forms of cell death (Horvitz 2003). However, its potential to study mechanisms underlying *patterned neurodegeneration* in response to human disease genes remains to be explored. This prompted us to ask whether APOE4 worms display a pattern of neuronal dysfunction, dysmorphia or degeneration beyond the HSN neurons as we previously reported (Sae-Lee et al., 2020). By day 4 (D4) of adulthood, APOE4 worms exhibit defects of varying severity in all seven tested behaviors. Because each behavior primarily relies on mostly distinct subsets of neurons (Fig. S1), this suggests a progressive, neuron-specific pattern analogous to the characteristic progression observed in human AD (Braak and Braak, 1991; Scahill et al., 2002; Fiford et al., 2018). The sensitivity of certain behaviors to decline merits follow up studies to investigate putative higher-vulnerability neurons (e.g., the ADE and PDE neurons required for swim-crawl gait transition; Vidal-Gadea et al., 2011; Fig. 1D). Certain behaviors were not altered by pan-neuronal APOE4 expression, perhaps indicating resilience to PTL-1-mediated degeneration or dysfunction. This idea is mirrored in human AD where brain regions such as the somatosensory cortex and brainstem are similarly spared from degeneration, even in advanced stages of AD (Rondina et al., 2018). Alternatively, apparent resistance to APOE4 may reflect neuronal circuit redundancy. In this vein, the subtle defects we observed in locomotion could reflect the functional redundancy of *C. elegans* motor neurons (Izquierdo and Beer, 2018; Fig. 1E,F). Interestingly, harsh-touch sensitivity subtly *increased* in APOE4 worms, hinting at neuronal hyperexcitability (Fig. 1H). In mouse, APOE4-driven hyperexcitability may result from degeneration of inhibitory neurons (Nuriel et al., 2017a). This aligns with fMRI studies of AD patients showing abnormally high activity in certain brain regions (Bookheimer et al., 2000; Tuovinen et al., 2020). Future studies could explore how APOE4 alters excitability of harsh touch neurons (e.g., the PVDs) and how this may correlate with APOE4-induced progressive dysmorphia of the elaborate PVD dendrite arborization (Jiang et al., 2025).

Axon dysmorphia, or axonopathy, is a hallmark of human AD characterized by neurite deterioration and general white matter abnormalities (Lee et al., 2016). Similar axon defects, sometimes without cell body loss, have been observed in human AD patients and mouse neurodegenerative disease models (Leroy et al., 2007; Yoshiyama et al., 2007; Crespo-Biel et al., 2014). Here, we identified APOE4-induced dysmorphia of the HSN proximal axon (Fig. 3). Our findings suggest this reflects a degenerative phenotype rather than a developmental defect because: HSN soma appear to migrate appropriately (Fig. S7), full axons project as expected in pre-adult L4 stage animals (Fig. S9), and the incidence and timing of abnormal axon morphology in D1 versus D3 adult APOE4 worms closely follow the HSN-associated egg laying defect (Figs. 1G, 3E). Like neurons in mammals, worm neurons have been shown to contain axonal domains with distinct structures and functions (Mizumoto and Shen, 2013; Eichel et al., 2022). Future studies should consider what molecular factors might render the proximal HSN axon vulnerable to dysmorphia. Overall, the tight correlation between a salient behavioral phenotype (bagging; Fig. 1G, 3G) and axon dysmorphia of an identified neuron (HSN; Fig. 3) provides a powerful system to study subcellular mechanisms of APOE4-induced axonopathy.

### Modeling degenerative mechanisms of tau in *C. elegans*

*C. elegans* has become increasingly recognized as a valuable *in vivo* model for studying tau pathology (Reviewed in Natale et al., 2020). Along with extracellular amyloid-β plaques, intracellular tau aggregation is a key pathological hallmark of AD (Kowall and Kosik, 1987; Kosik et al., 1988; Grunke-Iqbal et al., 1986; Lee and Trojanowski, 1992). In AD and other tauopathies, tau undergoes conformational changes that promote aggregation into tangles, driving neurodegeneration (Friedhoff et al., 1998a,b; Jeganathan et al., 2006; Jeganathan et al., 2008; Oakley et al., 2020). Unlike mammals, which have three broad tau-related gene families, the worm has but a single endogenous homolog: *ptl-1* (Sündermann et al., 2016; Goedert et al., 1996; Gordon et al., 2008). We found that deletion of *ptl-1* suppresses APOE4-induced defects in four out of seven tested behaviors—including HSN-associated bagging—*and* HSN axon morphological defects induced by APOE4. This parallels a previous report in a mouse AD model in which APOE4-induced dysfunction of specific interneurons was suppressed by knocking out endogenous murine tau (Andrews-Zwilling et al., 2010). Indeed, past immunohistochemical evidence suggested that APOE and tau co-localize – and possibly physically interact – in postmortem brain tissue from individuals with AD (Benzing and Mufson, 1995; Brecht et al., 2004; Zhou et al., 2006; Rohn et al., 2012). Further studies showed that APOE4 expression induces tau aggregates in cultured mouse neurons and worsens tau pathology in an AD mouse model (Huang et al., 2001; Seo et al., 2023; Koutsodendris et al., 2023 Shi et al., 2017). It will be interesting to see whether endogenous worm PTL-1 has similar pathological capacity detectable at the biochemical level.

Depletion of endogenous tau was reported to protect against neurodegeneration in an amyloid-β mouse model (Roberson et al., 2007). However, because *C. elegans* lacks an *APP* ortholog whose protein product is capable of generating amyloid-β fragments (Daigle and Li, 1993), the benefits of PTL-1 reduction likely operate independently of amyloid-β. Interestingly, a recent study found that *ptl-1* deletion partly mitigated age-dependent accumulation of pathogenic human tau upon its overexpression in worm neurons (Nunez et al., 2022). Thus, the APOE4 worm model of AD represents an Aβ-agnostic model to investigate how endogenous tau contributes to patterned degeneration downstream of APOE4.

How might worm tau contribute to degeneration in our APOE4 worm model? PTL-1 may contribute *cell-autonomously*. Notably, the pattern of behavioral decline roughly correlates with endogenous *ptl-1* expression levels in the primary neurons underlying each behavior (Fig. S1). This supports a model in which differences in baseline *ptl-1* expression set neuronal vulnerability, with higher-expressing neurons reaching a pathological threshold earlier than lower-expression neurons (Fig. S12). PTL-1 may also contribute *non-cell autonomously*. The TRNs contain the majority of PTL-1 in the worm nervous system (Taylor et al., 2021; Roux et al., 2023; Fig. S4 green bars). We found that depleting PTL-1 in the TRNs via genetic ablation, RNAi-mediated knockdown, and Cre-LoxP-mediated knockout all partially suppressed APOE4-induced dysfunction of HSN neurons. Each targeted manipulation roughly matched the magnitude of suppression achieved by systemic *ptl-1* deletion. This raises the intriguing possibility that, although *ptl-1* remains widely expressed, PTL-1 from the mere six TRNs is required non-cell autonomously for HSN dysfunction.

Tau spread represents a popular non-cell autonomous hypothesis to explain the link between tau pathology and spatiotemporal neurodegeneration in AD (Braak and Braak, 1991; Reviewed in Braak and Del Tredici, 2011; Frost et al., 2010). Studies with tauopathy mouse models have shown that pathological tau can propagate across synapses, starting from "seed" regions rich in Tau (and high in inter-neuronal connectivity) and spreading to distant brain regions (Clavaguera et al., 2009; Kfoury et al., 2012; Liu et al., 2012; de Calignon et al., 2012; Iba et al., 2013). Similarly, we hypothesize that pathological PTL-1 may spread from neuron to neuron over time in worm (Fig. S12). Although the six TRN soma are relatively distant from the two HSN soma (1/2-1/3 body length or ∼30-50 μm), PTL-1 is also enriched in TRN processes which pass nearby the HSNs (Fig. 4A2). Two TRNs (the PLM pair) also form chemical synapses with the HSNs (White et al., 1986). Future work could leverage synaptic transmission mutants to test whether this represents a site of PTL-1 spread (Miller et al., 1996; McColluch et al., 2017). If the hypothetical PTL-1 spread is synaptic, genetically perturbing TRN neurotransmission may phenocopy our targeted manipulations described above, suppressing HSN dysfunction. Because fluorescently tagged PTL-1 did not interfere with PTL-1-dependent degeneration of the HSNs (compare APOE4 strains with mNG tagged PTL-1 in Figs. 4,5 to non-tagged PTL-1 in Fig. 1G), our model may be used to directly visualize PTL-1 spread *in vivo*. With its simpler and fully mapped nervous system, *C. elegans* offers a simple model to investigate how endogenous tau propagates *in vivo* and contributes to APOE4-induced patterned neurodegeneration.

Deletion of worm tau did not equally protect against APOE4-induced behavioral dysfunction. We found that deletion of *ptl-1* suppresses APOE4-induced defects in swim-to-crawl switching and egg-laying (“bagging”) more strongly than other behaviors (Fig. 1D,G). Might inter-neuronal spread of tau explain this phenomenon? The synaptic connectivity of the neurons underlying these behaviors offers a possible explanation (White et al., 1986). Swim-to-crawl switching depends mainly on the dopaminergic neurons ADE and PDE, and egg-laying on the HSN neurons (Trent et al., 1983; Vidal-Gadea et al., 2014). Notably, the PTL-1-rich TRNs make direct chemical synapses onto these cells: AVM synapses onto ADE, and PLM synapses onto PDE and HSN. In contrast, *ptl-1* deletion does not suppress APOE4-induced abnormalities in short- or long-term locomotion or harsh-touch responses (Fig. 1E,F,H). Locomotion relies on many motor neurons, and harsh touch depends on a redundant sensory circuit including PVD, FLP, and touch neurons. Importantly, the TRNs do not directly synapse onto the DB and VB motor neurons that drive forward locomotion, nor onto the PVD or FLP neurons involved in harsh-touch sensing. Together, these connectivity patterns suggest a model in which PTL-1 produced in TRNs contributes to dysfunction in their postsynaptic partners (Fig. S12).

An alternate non-mutually exclusive hypothesis is that reduction of tau triggers organism-wide or neuroprotective effects. The idea that tau levels set a susceptibility threshold to many insults, not just tau-seeding pathology, may explain the broadly protective effects for deleting tau in rodent models. For instance, Roberson et al. (2007) found that halving or deleting endogenous tau rescued learning and memory as well as premature mortality in hAPP mice without changing Aβ levels or plaques but also reduced susceptibility to chemically induced seizures in otherwise wild-type backgrounds. Reducing tau also diminished spatial learning and memory deficits after mild repetitive traumatic brain injury and attenuated axonopathy and other histopathological changes in mice (Cheng et al, 2014). Systemic deletion of *ptl-1* reduces lifespan in *C. elegans* (Chew et al., 2013; Chew et al., 2014) and was shown to be necessary and sufficient in neurons for proper SKN-1-mediated response to environmental stressors (Chew et al., 2015). It remains untested whether (and in what tissues) fine-tuning of PTL-1 levels may confer general protection against environmental and/or neurotoxic stressors in young adult worms.

Regardless of how *ptl-1* contributes to degeneration, some APOE4-induced behavioral defects persist even when *ptl-1* is deleted. Therefore, APOE4 must also impair neuronal function through *ptl-1*–independent mechanisms. Future studies should investigate all the above possibilities.

## Materials and Methods

### Strain Maintenance

Unless otherwise noted, worms were maintained at room temperature (∼20 °C) on NGM plates seeded with OP50 bacteria as described (Brenner et al. 1974). Strains with extrachromosomal transgenes were propagated using selectable fluorescent reporters. Strains are listed in order of appearance in figures:

**Figures 1 and S2**

JPS1451 *vxEx1211[Pmyo-3::mCherry::UNC54UTR]*
RB809 *ptl-1(ok621) III*
JPS1453 *vxEx1213[Prab-3::ND18ApoE4::UNC54UTR + Pmyo-3::mCherry::UNC54UTR]*
JPS1455 *vxEx1215[Prab-3::ND18ApoE4::UNC54UTR + Pmyo-3::mCherry::UNC54UTR]; ptl-1(ok621) III*

**Figure 2**

JPS1451 *vxEx1211[Pmyo-3::mCherry::UNC54UTR]*
JPS1453 *vxEx1213[Prab-3::ND18ApoE4::UNC54UTR + Pmyo-3::mCherry::UNC54UTR]*

**Figures 3, S6, and S7**

JPS1666 *vxIs591[ptph-1::GFP::unc-54UTR]; vxEx1211[Pmyo-3::mCherry::UNC54UTR]*
JPS1664 *vxIs591[ptph-1::GFP::unc-54UTR]; vxEx1211[Pmyo-3::mCherry::UNC54UTR]; ptl-1(ok621) III*
JPS1667 *vxIs591[ptph-1::GFP::unc-54UTR]; vxEx1213[Prab-3::ND18ApoE4::UNC54UTR + Pmyo-3::mCherry::UNC54UTR]*
JPS1778 *vxIs591[ptph-1::GFP::unc-54UTR]; vxEx1213[Prab-3::ND18ApoE4::UNC54UTR + Pmyo-3::mCherry::UNC54UTR]; ptl-1(ok621) III*
JPS1779 *vxIs59[ptph-1::GFP::unc-54UTR]; vxEx1213[Prab-3::ND18ApoE4::UNC54UTR + Pmyo-3::mCherry::UNC54UTR]; ptl-1(ok621) III*

**Figure 4**

JPS1571 *vxIs591[ptph-1::GFP::unc-54UTR]*
JPS1828 *vxEx1511[Pmec-17::EGL-1::egl-1UTR]; ptl-1(pg73[PTL-1::mNeonGreen])*
TU253 *mec-4(u253) X*
JPS1500 *vxIs824[prab-3::ND18ApoE4::UNC54UTR, pmyo-2::mCherry::UNC54UTR]; ptl-1(pg73[PTL-1::mNeonGreen])*
JPS1839 *vxIs824[prab-3::ND18ApoE4::UNC54UTR, pmyo-2::mCherry::UNC54UTR]; ptl-1(pg73[PTL-1::mNeonGreen]); vxEx1511[Pmec-17::EGL-1::egl-1UTR]*
JPS1709 *vxIs824[prab-3::ND18ApoE4::UNC54UTR, pmyo-2::mCherry::UNC54UTR]; ptl-1(pg73[PTL-1::mNeonGreen]); mec-4(u253) X*

**Figure 5**

JPS1571 *vxIs591[ptph-1::GFP::unc-54UTR]*
GN655 *ptl-1(pg73[PTL-1::mNeonGreen])*
JPS1456 *ptl-1(pg73[PTL-1::mNeonGreen]); vxEx1216[Pmec-17::ptl-1 sense and antisense CDS + Punc-122::RFP]*
JPS1819 *ptl-1(syb8686 syb8597 pg73[LoxP::PTL-1::mNeonGreen::LoxP) III; hrtSi99[Pmec-17::Cre]*
JPS1872 *ptl-1(syb8686 syb8597 pg73) III; mirIs42[Pche-2::Cre + Plin-44::GFP]*
JPS1500 *vxIs824[prab-3::ND18ApoE4::UNC54UTR, pmyo-2::mCherry::UNC54UTR]; ptl-1(pg73[PTL-1::mNeonGreen])*
JPS1581 *vxIs824[prab-3::ND18ApoE4::UNC54UTR, pmyo-2::mCherry::UNC54UTR]; ptl-1(pg73[PTL-1::mNeonGreen]); vxEx1216[Pmec-17::ptl-1 sense and antisense CDS + Punc-122::RFP]*
JPS1582 *vxIs824[prab-3::ND18ApoE4::UNC54UTR, pmyo-2::mCherry::UNC54UTR]; ptl-1(pg73[PTL-1::mNeonGreen]); vxEx1216[Pmec-17::ptl-1 sense and antisense CDS + Punc-122::RFP]*
JPS1823 *vxIs824[prab-3::ND18ApoE4::UNC54UTR, pmyo-2::mCherry::UNC54UTR]; ptl-1(syb8686 syb8597 pg73[LoxP::PTL-1::mNeonGreen::LoxP) III; hrtSi99[Pmec-17::Cre]*
JPS1824 *vxIs824[prab-3::ND18ApoE4::UNC54UTR, pmyo-2::mCherry::UNC54UTR]; ptl-1(syb8686 syb8597 pg73[LoxP::PTL-1::mNeonGreen::LoxP) III; hrtSi99[Pmec-17::Cre]*
JPS1898 *vxEx1604[pRAB-3::mCherry::unc-54utr]; ptl-1(syb8686 syb8597 pg73[LoxP::PTL-1::mNeonGreen::LoxP) III; hrtSi99[Pmec-17::Cre]*
JPS1899 *vxEx1605[pRAB-3::mCherry::unc-54utr + Prab-3::ND18ApoE4::UNC54UTR]; ptl-1(syb8686 syb8597 pg73[LoxP::PTL-1::mNeonGreen::LoxP) III; hrtSi99[Pmec-17::Cre]*
JPS1897 *vxIs824[prab-3::ND18ApoE4::UNC54UTR, pmyo-2::mCherry::UNC54UTR];ptl-1(syb8686 syb8597 pg73) III; mirIs42[Pche-2::Cre + Plin-44::GFP]/+*

**Figure S8**

GN655 *ptl-1(pg73[PTL-1::mNeonGreen])*
JPS1500 *vxIs824[prab-3::ND18ApoE4::UNC54UTR, pmyo-2::mCherry::UNC54UTR]; ptl-1(pg73[PTL-1::mNeonGreen])*
JPS1828 *vxEx1511[Pmec-17::EGL-1::egl-1UTR]; ptl-1(pg73[PTL-1::mNeonGreen]) JPS1456 ptl-1(pg73[PTL-1::mNeonGreen]); vxEx1216[Pmec-17::ptl-1 sense and antisense CDS + Punc-122::RFP]*
JPS1819 *ptl-1(syb8686 syb8597 pg73[LoxP::PTL-1::mNeonGreen::LoxP) III; hrtSi99[Pmec-17::Cre]*

### Generation of strains

The 1933 bp deletion in *ptl-1* allele *ok621* was confirmed by PCR using the following primers (written in 5’-3’ orientation):

Forward: CTGGAAATTTGTTGGGCAGT
Reverse: TGAACCGAAGCCTAAACCAG

We sought to independently confirm the effect of pan-neuronal APOE4 on egg laying as well as to test other behaviors with new extrachromosomal transgenic strains that carry a new reporter that minimized effects on multiple behaviors. We transformed wild-type N2 with 20 ng/μL of the APOE4 plasmid pPS90 (Sae-Lee et al., 2020) along with 10 ng/μL of the pCFJ104 mCherry body wall reporter (Frøkjaer-Jensen et al., 2008) to generate the new extrachromosomal strain JPS1453. In parallel, we transformed the *ptl-1*(*ok621*) mutant strain RB809 with the same DNA mixture yielding the APOE4;Δ*ptl-1* strain JPS1455. We also generated control strain JPS1451 that carries the mCherry reporter alone.

To visualize HSN axons, we first used the GFP reporter strain JPS1571. This strain was crossed with JPS1451 and JPS1453 to yield the control and APOE4 strains JPS1666 and JPS1667, respectively. JPS1666 was then crossed with RB809 to yield the *Ptph-1::gfp*;Δ*ptl-1* strain JPS1664. JPS1667 was crossed with RB809 to yield APOE4;Δ*ptl-1*, from which two independent isolates were obtained, JPS1778 and JPS1779, to partly control for genetic background (Fig. 3). In our independent reporter lines, we transformed strain JPS1819 with 20 ng/uL of the mCherry reporter plasmid pGH8 without or with 40 ng/uL APOE4 plasmid pPS90 to yield strains JPS1898 and JPS1899 (Fig. 5B3).

### Cell-specific gene knockdown

To knock down *ptl-1* specifically in the TRNs, we followed previous methods by Esposito et al. (See Fig. 1 of Esposito et al., 2007). The *mec-17* promoter (1535 bp) was fused with either sense or antisense of *ptl-1* coding region (820 bp) to generate separate DNA fragments using the following primers (written in 5’-3’ orientation):

A

Forward: GGCTTGAATAATCCCTATTATAGCC
Reverse: GAAATTACCTGAAATTTCGTGGGG

B

Forward: GAGAAAACGGCCAATTGACGC
Reverse: GGCTATAATAGGGATTATTCAAGCCGATCGAATCGTCTCACAACTG

C

Forward: GAGAAAACGGCCAATTGACGC
Reverse: CCCCACGAAATTTCAGGTAATTTCGATCGAATCGTCTCACAACTG

D

Forward: CGTCGCGCTAACAGTTCTAG
Reverse: CCAAAGGTTAACGCCAAATTTG

E

Forward: CGTCGCGCTAACAGTTCTAG
Reverse: CAAGTGAAAGCGTGGTCCGATC

Fragments D and E were mixed in equimolar amounts, along with a selectable RFP coelomocyte marker (Addgene Plasmid #8938), to a final concentration of 80 ng/μL. This DNA mix was transformed into the *ptl-1*::*mNeonGreen* strain GN655 (PTL-1::mNG; Krieg et al., 2017). The resultant strain, JPS1456, expresses *ptl-1* dsRNA fragments in the TRNs. JPS1456 was crossed into JPS1500, which carries an integrated pan-neuronal APOE4 transgene; this yielded APOE4;TRNs::*ptl-1* KD from which two independent isolates were obtained, JPS1581 and JPS1582, to partly control for genetic background.

### Cell-specific gene knockout

To knock out *ptl-1* in the TRNs, we commissioned SUNY Biotech to generate the strain PHX8686 which harbors *LoxP* sites flanking the *ptl-1* genetic locus in the strain GN655. PHX8686 was crossed into MSB1167, a transgenic line in which Cre recombinase is expressed under control of the *mec-17* promoter (Harterink et al., 2018). This yielded the TRNs::*ptl-1* KO strain JPS1819. The presence of PTL-1::mNG in the TRNs was determined with epifluorescent microscopy. Lastly, JPS1819 was crossed into JPS1500 to yield APOE4;TRNs::*ptl-1* KO, from which two independent isolates were obtained, JPS1823 and JPS1824, to partly control for genetic background.

Similarly, knockout of *ptl-1* in the *che-2*-expressing subset of sensory neurons (CHEs) was achieved by crossing PHX8686 into FX16644, a transgenic line in which Cre recombinase is expressed under control of the *che-2* promoter (National BioResource Project, Japan). This yielded the CHEs::*ptl-1* KO strain JPS1872. However, subsequent crossing into JPS1500 resulted in synthetic lethality in animals homozygous for the CHEs-specific Cre transgene. Thus, heterozygotes were selected for egg laying behavior assays.

### Cell-specific ablation

To genetically ablate the TRNs, we used the TRN-specific *mec-17* promoter to express *egl-1* in an extrachromosomal array in strain JPS1828 (Wu et al., 2025). This strain was crossed into JPS1500, a strain carrying an integrated pan-neuronal APOE4 transgene and a *Punc-122*::*gfp* reporter, to yield strain JPS1839. Additionally, we crossed JPS1500 into TU253, which carries a *mec-4* loss-of-function mutation (O’Hagan et al., 2004), to yield the APOE4;*mec-4*(*lf*) strain JPS1709.

### Sterilization

To inspect HSN neuron health without confounding effects of the bagging phenotype, worms were sterilized as previously described (Mitchell et al., 1979). In brief, seeded plates were pretreated with 600 mL of 0.8 mM 5’-fluorodeoxyuridine (FUdR; FUdR plates). Mid- to late-stage L4 stage worms were transferred onto FUDR plates and moved to fresh FUdR plates every 1-2 days as needed.

### Behavioral assays

Behavioral assays were performed on either 60 mm (medium) or 35 mm (small) NGM plates, seeded or not with OP50 bacteria as described per assay. All assays were performed at room temperature (20° C) unless stated otherwise. Assays began with day 1 adults (D1), 24 hours after initial incubation on FUDR plates (if applicable), until a significant difference between control and APOE4 worms was observed. Completely immobile worms or those which escaped from the side of the plate were omitted from assays and final quantifications.

### Gentle touch

Methods adapted from Chalfie and Sulston (1981). In brief, 20-30 worms were moved to medium NGM-OP50 plates. To test gentle touch sensitivity, an eyelash hair, affixed by its shaft to a pipette tip, was used to deliver mechanical stimuli to worms. Gentle strokes were applied to the posterior and anterior of the animal. Six paired anterior and posterior stimuli were applied, with an interstimulus interval of approximately 1.5 seconds to avoid habituation. Gentle touch sensitivity was quantified per worm as the fraction of responses to stimuli out of 12.

### Harsh touch

Methods adapted from Li et al. (2011). In brief, 20-30 worms were moved to medium NGM-OP50 plates. To test harsh gentle touch sensitivity, a single prod was delivered to the mid-body using a platinum pick. Five technical replicates were performed per worm per strain. Harsh touch sensitivity was quantified per replicate as the proportion of responsive worms.

### Pharyngeal pumping

Methods adapted from Avery and Shtonda (2003). In brief, 30-40 worms were first moved to medium NGM-OP50 plates. Worms were allowed to roam for at least 30 minutes. Individual animals were then tracked for 20 seconds and pharyngeal bulb contractions were manually tallied with a handheld click counter. Pumping rate was quantified per worm as pumps per minute.

### Bagging

Methods adapted from Sae-Lee et al. (2020). In brief, at least 30-40 L4-stage worms were moved to medium NGM-OP50 plates. Worms were scored daily for the internal hatching or “bagging” phenotype based on presence or absence of several late-stage eggs and/or hatched larvae within the uterus; this is often accompanied by general lethargy plus distension of the worm midbody as previously described (Trent et al., 1983). Bagging worms were tallied and discarded while non-bagging worms were transferred to a fresh medium NGM-OP50 plate. Worms were scored in this fashion from D1 to D4 of adulthood. Bagging incidence was quantified per replicate as the cumulative percentage of bagging worms summed across D1-4 out of the original total.

### Long-term locomotion

Methods adapted from Nordquist et al. (2018). Single worms were moved to the center of a small NGM plate uniformly seeded with OP50 such that the lawn covered the entirety of the NGM surface. Animals were incubated on these assay plates overnight at 22°C. Groups of at least 8 worms per genotype were tested on two separate days as also described by Nordquist et al. After 18 hours, the worms were removed from their plates and a transparent grid guide was used to determine the number of ∼5 mm^2^ regions (32 in total) traversed on the assay plate. Regions encompassing parts of the agar that split or pulled away from the plastic of the plate were omitted from the total. Long-term locomotion was quantified per worm as the proportion of traversed regions on the assay plate out of the total.

### Short-term locomotion

Methods adapted from Topalidou et al. (2017). In brief, 8-10 worms were moved to the center of a medium NGM-OP50 plate that was uniformly seeded such that the bacterial lawn extended across the entire medium surface. After 10 minutes, worms were removed and their final positions marked on the plate. Groups of least 15 worms per genotype were tested on two separate days as previously described (Nordquist et al., 2018). Radially dispersed distance was recorded. Short-term locomotion was quantified per worm as the distance (cm) from the center to final positions.

### Swim-to-crawl gait transition

Methods adapted from Vidal-Gadea et al. (2011). Before testing, 25-40 worms were moved to a medium NGM plate and allowed to roam for ∼15 minutes so excess OP50 could be removed. Meanwhile, assay plates were prepared by first drawing a 0.5 cm radius border on the plastic behind its center. Next, 10 uL of liquid NGM was pipetted to the very center of the assay plate surface. Worms were then picked to the center of the puddle on the assay plate and a 15-minute timer was started. As the puddle dried, swimming worms began to escape to the solid surface, transition to a crawling gait, and disperse away from the puddle. Gait transition was quantified per replicate as the proportion of worms that escaped and crawled past the 0.5 cm border in under 15 minutes.

### Microscopy

Epifluorescent microscopy was performed on an Olympus IX51 inverted microscope with an Olympus UPlanFL N 40X/0.75 NA objective and X-Cite FIRE LED Illuminator (Excelitas Technologies Corp.). Images were captured using a Retiga 2000R CCD camera (QImaging) and QCapture Pro 6.0 software.

Worms were sterilized with FUDR as described above to prevent internally hatching larvae that may affect HSN morphology. Worms were then mounted on 2% agarose pads and immobilized using 50 mM sodium azide. All imaging per pad took place within 30 minutes of mounting.

The same fluorescence intensity settings were used to image all HSN soma and proximal axons via a GFP reporter. Importantly, imaging was restricted to worms that lay in a strictly lateral orientation (i.e., on the right or left side with minimal twisting about the long axis of the animal). This ensured both the soma and proximal axon of a given HSN neuron were in approximately the same focal plane. We determined the orientation of mounted worms using the fluorescent co-injection marker. This marker labels body wall and vulval musculature with mCherry, visualizable in the red channel. In single focal planes, the labeled vulval muscles either appear as an x-shape from a ventral viewpoint or a v-shape if the animal laid directly on its right or left side as described above (See also Fig. 1 of Collins and Koelle, 2020). Incidentally, the HSN GFP reporter also labels the VC4 and VC5 motor neurons which are visible in the same focal plane. These guides helped restrict our microscopy to animals in the desired orientation described above.

After imaging, morphometrics of HSN soma and proximal axons were calculated in ImageJ (Schneider et al., 2012). Positions of HSN soma were determined by measuring the lengths of the vector from the soma to the ventral nerve cord (HSN-VNC, met at a perpendicular) and the vector to the vulval area in between VCs4,5 (HSN-vulva). These measurements were also collapsed into a single metric as previously described (Asakura et al., 2007). This metric, represented by θ, is the angle in degrees between the HSN-vulva (hypotenuse) and HSN-VNC (adjacent) vectors (i.e., arccosine of their ratio).

HSN proximal axon lengths were calculated with the NeuronJ plugin for ImageJ using default settings (Meijering et al., 2004). Traces were made by following the proximal axon from anterior face of the HSN soma to the point at which the axon exits the focal plane ventrally in the vulval area. Ectopic sprouts from the soma or proximal axon were excluded from tracing. In the rare (<5% frequency) cases where ectopic sprouts were present on the proximal axon, the shortest ventral- and vulval-directed course was traced for all strains imaged.

Full HSN axons were imaged in mid- to late-L4 stage worms when full projection to the head region is typically complete (Garriga et al., 1993). Worms with very dim GFP fluorescence were excluded from scoring, as it was ambiguous whether axons failed to project or the reporter transgene was not yet highly enough expressed.

To capture images of PTL-1::mNG strains, an increased exposure time and intensity setting were required compared to HSN GFP reporter strains. Because some TRN processes span nearly the entire animal length, we chose to focus on soma to quantify PTL-1::mNG signal in TRNs::*ptl-1* KD strain JPS1456 and control strain GN655 (See Fig. 4). Depending on animal orientation, bilateral TRN soma were identifiable based on the naturally asymmetric positioning of the AVM and PVM class neurons (Arnold et al., 2020). The VCs4,5 neurons were almost always visualizable regardless of orientation, making them ideal for internal controls. Fluorescence intensity was quantified in ImageJ in day 1 adults.

In the TRNs::*ptl-1* KO strain JPS1819, *LoxP*-flanked *ptl-1* is presumably excised by Cre recombinase whose expression is driven by the *mec-17* promoter (Harterink et al., 2018). Although *mec-17* expression begins embryonically in ALM and PLM class TRNs (Roux et al., 2023), residual PTL-1::mNG is still present by at least L4 stage of development; PTL-1::mNG is qualitatively fainter in those cells compared to like-aged control animals (GN655 strain). Thus, selection for animals with lacking PTL-1::mNG in PLM neurons was conducted in at least day 1 adults to avoid ambiguity. This selection strategy was also employed in the TRNs::EGL-1 strain JPS1828 to identify animals with genetically ablated TRNs.

### Western blotting

Worms were washed twice with 5% sucrose and twice with 50 mM NaCl before being flash frozen with 5μl glass beads. For worm lysis, 2% PMSF was added to 2x Laemmli buffer. This buffer was then addedto the frozen samples to achieve a 1x concentration. Finally, 5% β-mercaptoethanol (BME) was added to the samples, followed by bead beating for 30 seconds. Samples were spun down through centrifugation for 2 minutes at 15,000 g. The samples were then boiled a t 95℃ for 5 minutes. An additional 2-minute centrifugation was performed to clear debris prior to measuring protein concentration with Qubit Protein Assay Kit (Thermo Scientific, #Q33211).

20 μg of protein per sample was loaded on Invitrogen™ NuPAGE™ Bis-Tris Mini Protein Gels (4–12%, 1.0–1.5 mm, Invitrogen, #NP0335) and separated at 80-120 V in 1X MES SDS running buffer. Proteins were transferred onto a PVDF membrane (Thermo Scientific, #88518) at 30V for 1 hour.

Membranes were blocked in 1X TBS Tween 20 Buffer (Thermo Scientific, #28360) containing 5% milk. Flag tag and actin were probed with primary antibodies: Monoclonal ANTI-FLAG® M2 antibody produced in mouse (1:2,000, Sigma-Aldrich, #F3165) and Anti-Actin Mouse Monoclonal Antibody [clone: C4] (1:500, MP Biomedicals, #08691001), respectively. A secondary antibody, Goat anti-Mouse IgG (H+L) Superclonal™ Secondary Antibody, Alexa Fluor™ 488 (1:4,000, Invitrogen, #A28175) was used after flag and actin probing. Finally, an Amersham™ Typhoon™ RGB was utilized for visualization.

### Statistical analysis

All data are presented as the mean ± SEM. Plotted data points represent individuals or the average across a single replicate as indicated in figures. Total sample size (N) is indicated in figure legends. Behavioral assays and microscopy scoring were performed blind to genotype. Planned χ^2^ tests of independence were performed for statistical comparisons of gait transition (escaped vs. not escaped), bagging (yes vs. no), and HSN axon scoring (normal vs. abnormal) in Excel 2022 (365 MSO, Version 2510 Build 16.0.19328.20190, 64-bit). Touch, pumping, and locomotion behaviors, HSN axon length, HSN soma position, and fluorescence intensity were compared with planned, Student t-tests or post hoc comparisons following two-way ANOVA tests (SPSS, v29.0.2.0, IBM).

## Acknowledgements

We would like to thank Drs. Miriam B. Goodman and Michael Krieg for sharing strains GN655 and MSB1167, respectively, and SUNY Biotech for engineering PHX8686. Some strains were also provided by the *Caenorhabditis* Genetics Center, which is funded by the NIH (P40 OD010440),. We also thank Dr. Lotti Brose and Anna Krauss for preliminary experiments with APOE4; Δ*ptl-1* animals.

## Footnotes

### Author contributions

Conceptualization: J.T.P., E.A.C.; Methodology: E.A.C., Z.W., J.T.P.; Investigation: E.A.C., Z.W., B.M.B., C.J.W., E.S.-C., J.T.P.; Writing – original draft: E.A.C., J.T.P.; Writing – review and editing: E.A.C., Z.W., C.J.W., E.S.-C., J.T.P.; Funding acquisition: E.S.-C., J.T.P.

### Data availability

All relevant data can be found either within the article or in its supplementary information section. Raw data is available upon request.

