## Supplemental Info for "Loss of endogenous tau suppresses APOE4-induced patterned behavioral decline and axon dysmorphia in a *C. elegans* model of Alzheimer’s disease"

#### Supplemental Material

**Fig. S1.**

**Behavior-associated neurons vary in number, how much *ptl-1* they express, and timing of potential APOE4-induced dysfunction**

Behaviors became defective or abnormal at different ages (grouped by age of onset in dashed ovals). We speculate that this timing may relate to the amount of *ptl-1* expressed across behavior-associated neurons, depicted by the grey circle size nearby each behavior.

**Fig. S2.**

**Continued assessment of harsh touch out to adult day 6 (D6)**

Compared to control, APOE4 ($p<0.001$) and APOE4;Δ*ptl-1* ($p<0.01$) animals are more sensitive to harsh touch beginning in D3 adults. N=25 for control, N=26 for APOE4, and N=31 for APOE4;Δ*ptl-1*. To explore this unexpected increase in sensitivity, animals were also tested up to D6 age adults. Compared to D4 control, the increased sensitivity of APOE4 animals remains ($p<0.01$; N=27 for control and N=20 for APOE4), but no statistically significant difference was observed at D5 (N=20 for control and N=20 for APOE4) into D6 (N=12 for control and N=20 for APOE4).

All data are represented as mean ± SEM. All statistical comparisons were made using planned tests after two-way ANOVA, where # or @ indicate significant differences between control and APOE4 or control and APOE4;Δ*ptl-1* groups, respectively. N is the total sample size representing all animals across replicates.

**Fig. S3.**

**Replotted: APOE4-induced behavioral decline may be the result of developmental and age-related defects**

Replotting of Figure 2A-C to compare strains and days in one graph. (A) Assays of gentle touch sensitivity. (B) Assays of pumping.

**Fig. S4**

**Nervous system *ptl-1* expression**

Bar plot showing cell-specific RNA sequencing data acquired from CeNGEN data set (Taylor et al., 2021). Green bars represent the touch receptor neurons (TRNs), blue bars represent egg-laying associated neurons (VCs4,5 and HSN), and yellow bars represent the *che-2*-expressing sensory neurons (CHEs).

**Fig. S5.**

**Disease-relevant tau domains are conserved from human to *C. elegans***

Canonical microtubule binding repeats (R1-4) of human tau are marked above the alignment using double-sided arrows. Enclosed by red boxes are protein domains shown to interact with each other within tau aggregates across tauopathies (Reviewed in Oakley et al., 2020). Yellow arrowheads indicate residues whose mutation has been empirically shown to promote or inhibit aggregate formation of human tau (McDermott et al., 1996; Strang et al., 2019); these residues are also conserved in worm PTL-1. Alignment generated with Clustal Omega using publicly available primary sequences for endogenous tau protein in human and worm (The UniProt Consortium; Wormbase). Note that full length tau is not depicted to emphasize the R domains.

**Fig. S6.**

**Morphometrics of the HSN soma are inconsistent correlates of APOE4-induced dysfunction**

(A) Descriptions of the soma morphometrics quantified in ImageJ with the goal of identifying characteristics which we hypothesized would correlate with our behavioral data (See Fig. 1G).

(B) Quantification of HSN soma area. While APOE4 animals exhibited larger soma on average compared to control, so did APOE4 and APOE4;Δ*ptl-1* groups.

(C) The circularity metric of the HSN soma was not obviously reflective of HSN dysfunction.

(D-F) Quantification of average GFP reporter brightness in the HSN soma.

(D) On average, HSN soma in APOE4 animals were dimmer, qualitatively consistent with our previous observations (Sae-Lee et al., 2020). However, APOE4;Δ*ptl-1* animals exhibit even dimmer HSN soma than APOE4.

(E,F) Quantifications of pixel intensity distribution. APOE4 reporter distributions appear, on average, to be more tail-skewed and exhibit more positive kurtosis compared to control. This could indicate non-uniform filling of GFP in the cytoplasm, but how this reflects HSN health and function is unclear.

All data are represented as mean ± SEM. N is the total sample size representing all animals. N=19 for control, N=9 Δ*ptl-1*, N=18 for APOE4, N=12 for APOE4;Δ*ptl-1* (left), and N=12 for APOE4;Δ*ptl-1* (right).

**Fig. S7.**

**Normal HSN axons in D1 adults and HSN soma positions in D3 adults**

(A) Cartoon of the worm midbody, centered about the vulva area, depicting: the HSN proximal axon, the VC4,5 neurons, vulval musculature, and ventral nerve cord (VNC)

(B) Representative live images of both normal HSN axon morphologies in D1 adult control animals. The vulva area is also marked with a white arrowhead for anatomical reference. All scale bars are 20 µm.

(C) Lengths of vectors measured from either soma to VNC met at a perpendicular or soma to vulva area as indicated by arrowheads of respective dashed lines (left). Consolidated metric of HSN soma position calculated as the angle θ, in degrees, formed between the vectors described above (right). Quantification of HSN soma position shows no significant difference between control and APOE4. N=17 for control,

N=9 for Δ*ptl-1*, N=16 for APOE4, N=10 for APOE4;Δ*ptl-1* (left), and N=11 for APOE4;Δ*ptl-1* (right). Statistical comparisons were made using planned comparisons after two-way ANOVAs

**Fig. S8.**

**Replotted Figure 3: Morphometrics of the HSN soma are inconsistent correlates of APOE4-induced dysfunction**

Replotted Figure 3C-F to compare strains and days in one graph. (A) Percent of abnormal HSN axon shape. (B) Length of proximal HSN axon.

**Fig. S9.**

**HSN axons projection is normal at L4 developmental stage in APOE4 worms**

(A) Cartoon depicting the midbody and head areas of the worm body through which the HSN axon typically projects.

(B) Quantification of animals with normally projecting axons through the ventral nerve cord (VNC) into the head in control (N=28) and APOE4 (N=22) worms.

(C,D) Representative live images of HSN (soma in dashed circle) axon projections in the VNC (arrowheads) and head (arrow) from animals in (C) lateral or (D) ventral orientations. All scale bars are 20 µm.

**Fig. S10.**

**Confirmation of PLM neuron cell body presence or absence using Differential Interference Contrast (DIC) and fluorescent microscopy.** Representative DIC and fluorescent micrographs of tail of different strains. (Top) Soma of PLM touch neuron in the tail is visible with DIC optics and mNeonGreen fluorescently tagged PTL-1 in control worm. (Middle) PLM soma is present but the mNG tagged PTL-1 is absent in the TRNs::ptl-1 worm. (Bottom) PLM soma and mNG tagged PTL-1 are both absent in the TRNs::EGL-1 worm in which the TRNs are genetically ablated.

**Fig. S11.**

**Whole-animal PTL-1 content from control strains with PTL-1 depleted in the TRNs**

(left) Representative Western blot to detect PTL-1::mNG. The expected band of ~76kDa was detected by probing for the FLAG epitope engineered into this tag (Krieg et al., 2017). (right) Western blot quantification of FLAG bands normalized to actin bands in *ptl-1*::*mNG* strains revealed variably levels of intensity for control (strain GN655) and the

TRNs::EGL-1 (Genetically ablated TRNs), TRNs::*ptl-1* KD, TRNs::*ptl-1* KO strains used in Figs. 4 and 5.

**Fig. S12.**
**Working model: Neuronal dysfunction and dysmorphy relates to different intrinsic levels and interneuron spread of worm tau.** In the control worm, HSN and PLM neurons function and appear normal when their levels of worm tau (PTL-1) are below a susceptibility threshold. In the APOE4 worm, APOE4 induces PTL-1 levels to rise above a pathological susceptibility threshold due to 1) accumulation within a neuron, and/or 2) synaptic spread. In this model, the PLM neuron becomes dysfunctional before the HSN neuron because its intrinsic levels of PTL-1 are higher. HSN neuron becomes dysfunctional and dysmorphic next because of pathological spread and accumulation.

Fig. S1

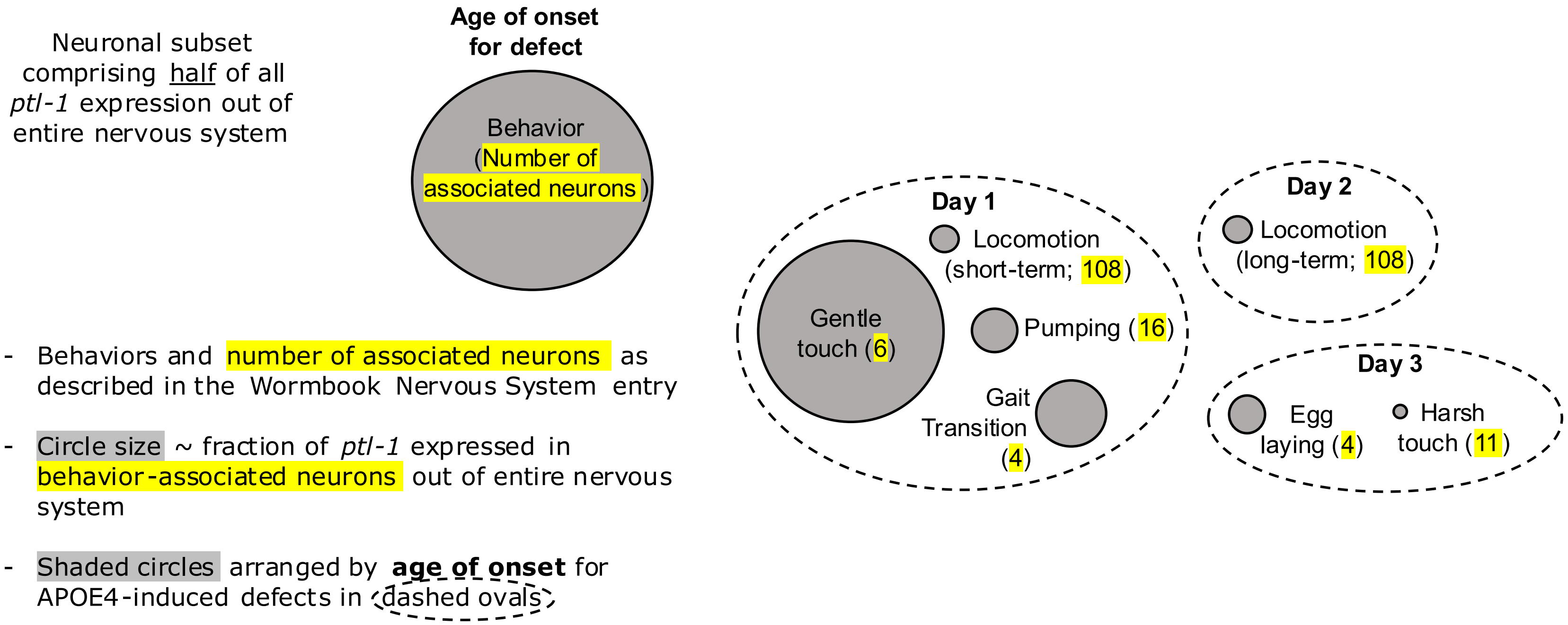
Neuronal subset comprising half of all ptl-1 expression out of entire nervous system
Age of onset for defect
Behavior (Number of associated neurons)
- Behaviors and number of associated neurons as described in the Wormbook Nervous System entry
- Circle size ~ fraction of ptl-1 expressed in behavior-associated neurons out of entire nervous system
- Shaded circles arranged by age of onset for APOE4-induced defects in dashed ovals
Day 1
Locomotion (short-term; 108)
Gentle touch (6)
Pumping (16)
Gait Transition (4)
Day 2
Locomotion (long-term; 108)
Day 3
Egg laying (4)
Harsh touch (11)

### Fig. S2

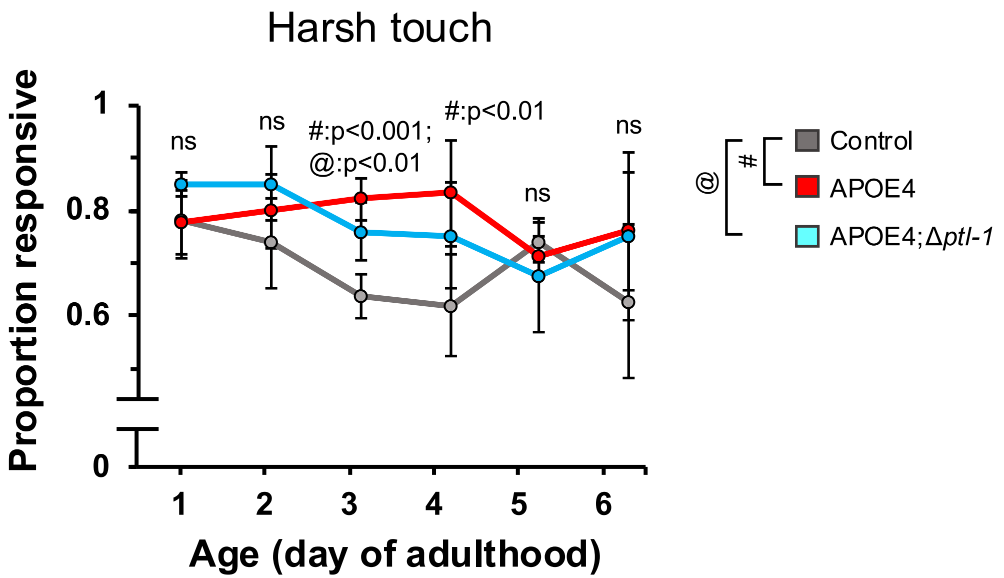

Harsh touch
Proportion responsive
1
0.8
0.6
0
ns
ns
#:p<0.001;
@:p<0.01
#:p<0.01
ns
ns
1
2
3
4
5
6
Age (day of adulthood)
@
#
Control
APOE4
APOE4;Δptl-1

Fig. S3

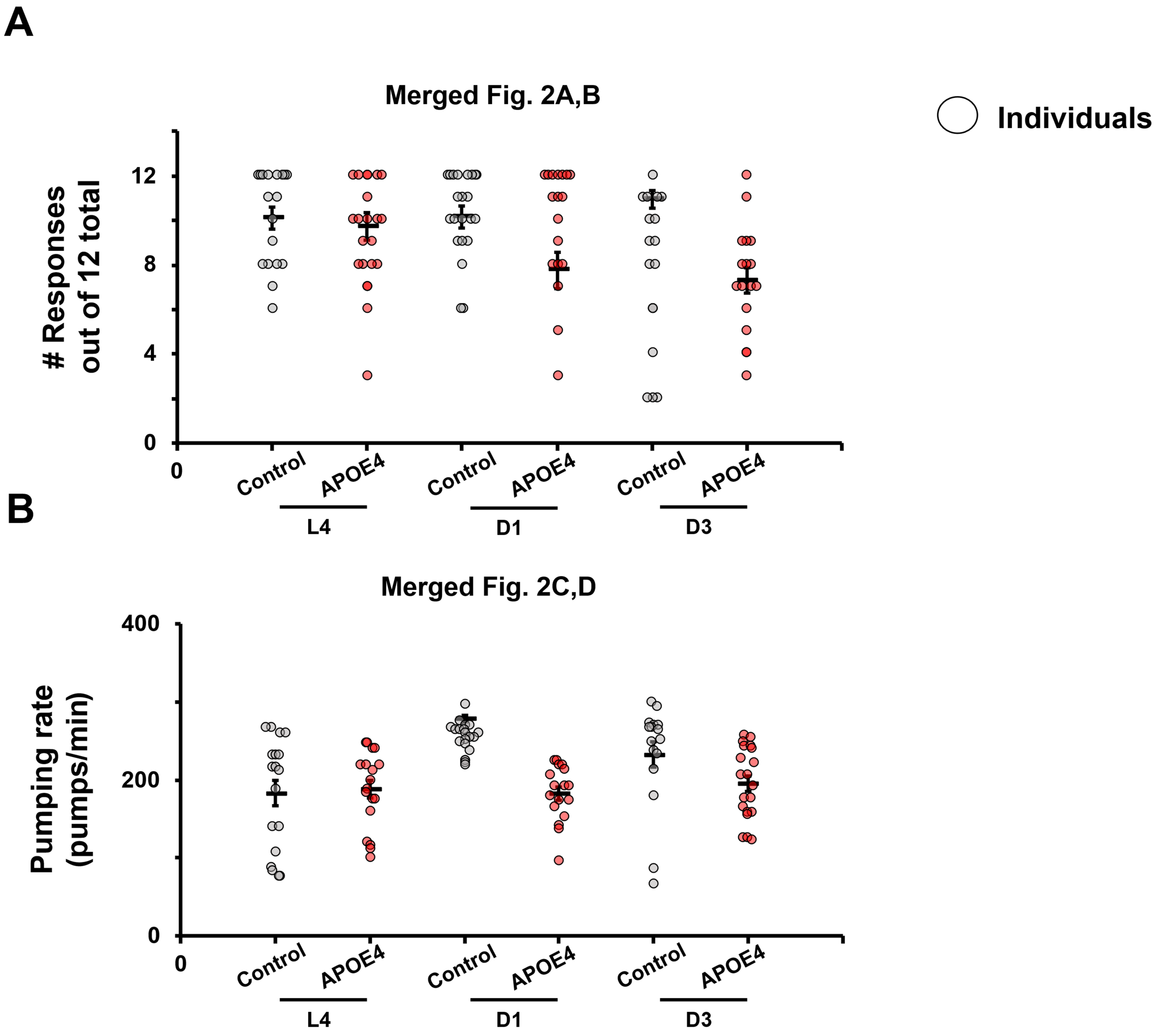
A
Merged Fig. 2A,B
### Responses
out of 12 total
0
4
8
12
0
Control
APOE4
Control
APOE4
Control
APOE4
L4
D1
D3
B
Merged Fig. 2C,D
Pumping rate
(pumps/min)
0
200
400
0
Control
APOE4
Control
APOE4
Control
APOE4
L4
D1
D3
Individuals

#### Fig. S4

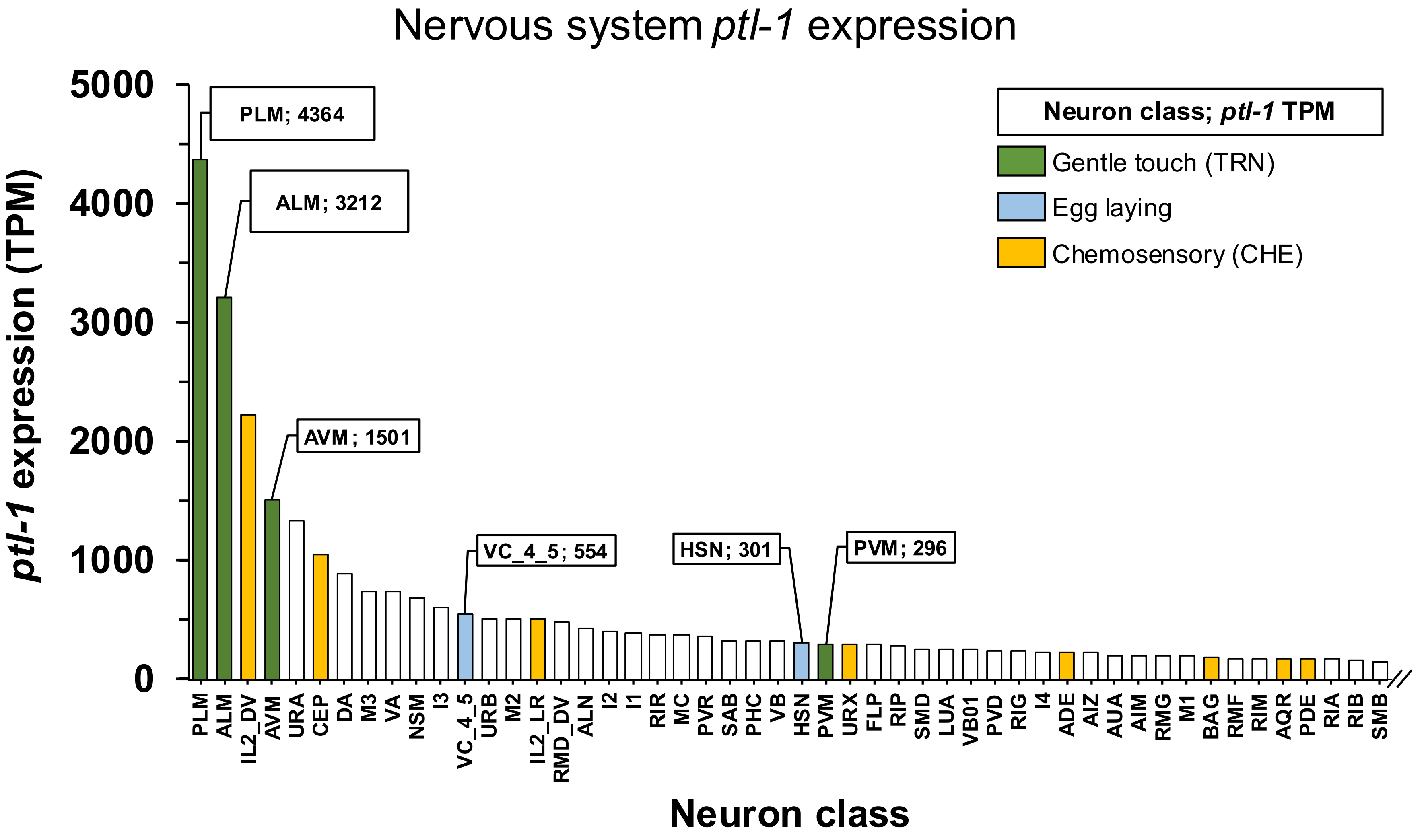

Nervous system ptl-1 expression
ptl-1 expression (TPM)
5000
4000
3000
2000
1000
0
PLM; 4364
ALM; 3212
AVM; 1501
VC_4_5; 554
HSN; 301
PVM; 296
Neuron class; ptl-1 TPM
Gentle touch (TRN)
Egg laying
Chemosensory (CHE)
PLM
ALM
IL2_DV
AVM
URA
CEP
DA
M3
VA
NSM
I3
VC_4_5
URB
M2
IL2_LR
RMD_DV
ALN
I2
I1
RIR
MC
PVR
SAB
PHC
VB
HSN
PVM
URX
FLP
RIP
SMD
LUA
VB01
PVD
RIG
I4
ADE
AIZ
AUA
AIM
RMG
M1
BAG
RMF
RIM
AQR
PDE
RIA
RIB
SMB
Neuron class

### Fig. S5

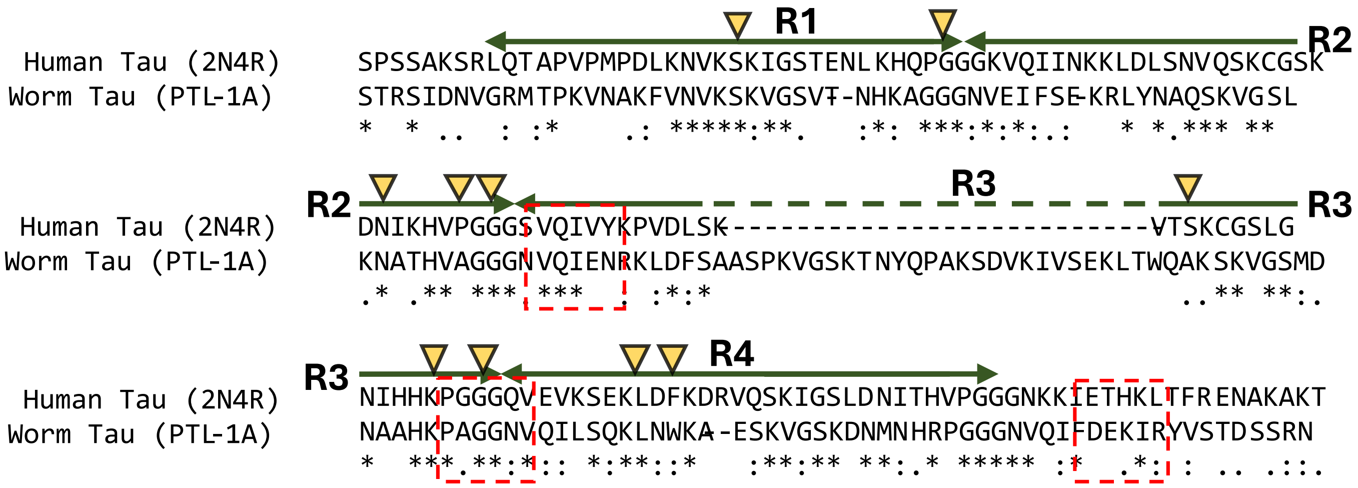
R1
R2
R3
R4
Human Tau (2N4R)
Worm Tau (PTL-1A)
SPSSAKSRLQTAPVPMPDLKNVKSKIGSTENLKHQPGGGKVQIINKKLDLSNVQSKCGSK
STRSIDNVGRMTPKVNAKFVNVKSKVGSVT-NHKAGGGNVEIFSE-KRLYNAQSKVGSL
DNIKHVPGGGSVQIVYKPVDLSK-------------------------VTSKCGSLG
KNATHVAGGGNVQIENRKLDFSAASPKVGSKTNYQPAKSDVKIVSEKLTWQAKSKVGSMD
NIHHKPGGGQVEVKSEKLDFKDRVQSKIGSLDNITHVPGGGNKKIETHKLTFRENAKAKT
NAAHKPAGGNVQILSQKLNWKA--ESKVGSKDNMNHRPGGGNVQIFDEKIRYVSTDSSRN

Fig. S6

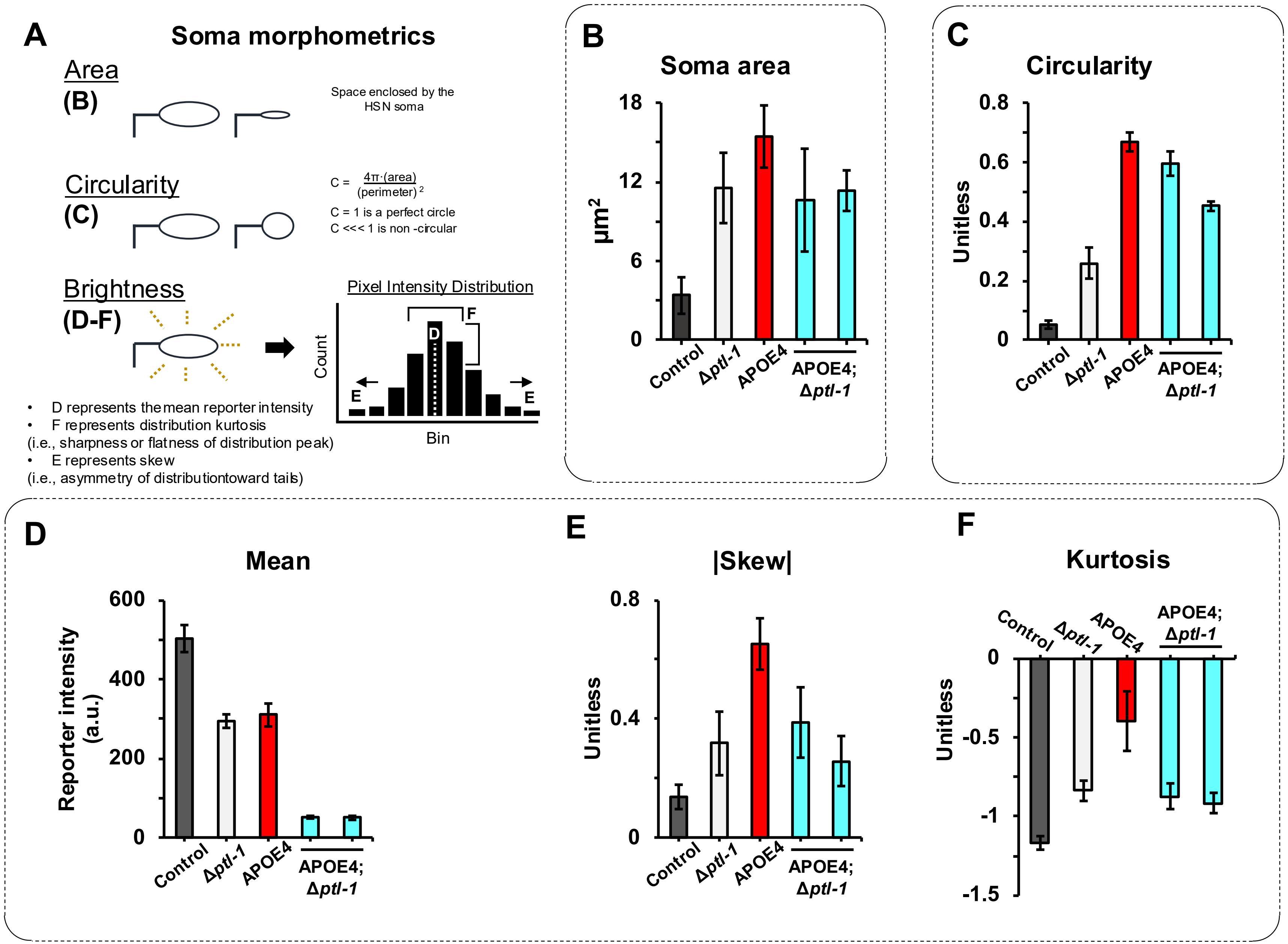
A
Soma morphometrics
Area
(B)
Space enclosed by the HSN soma
Circularity
(C)
C = 4π·(area) / (perimeter)²
C = 1 is a perfect circle
C <<< 1 is non -circular
Brightness
(D-F)
Pixel Intensity Distribution
Count
Bin
D
E
F
• D represents the mean reporter intensity
• F represents distribution kurtosis (i.e., sharpness or flatness of distribution peak)
• E represents skew (i.e., asymmetry of distribution toward tails)
B
Soma area
μm²
18
12
6
0
Control
Δptl-1
APOE4
APOE4; Δptl-1
C
Circularity
Unitless
0.8
0.6
0.4
0.2
0
D
Mean
Reporter intensity (a.u.)
600
400
200
0
E
|Skew|
Unitless
0.8
0.4
0
F
Kurtosis
Unitless
0
-0.5
-1
-1.5

Fig. S7

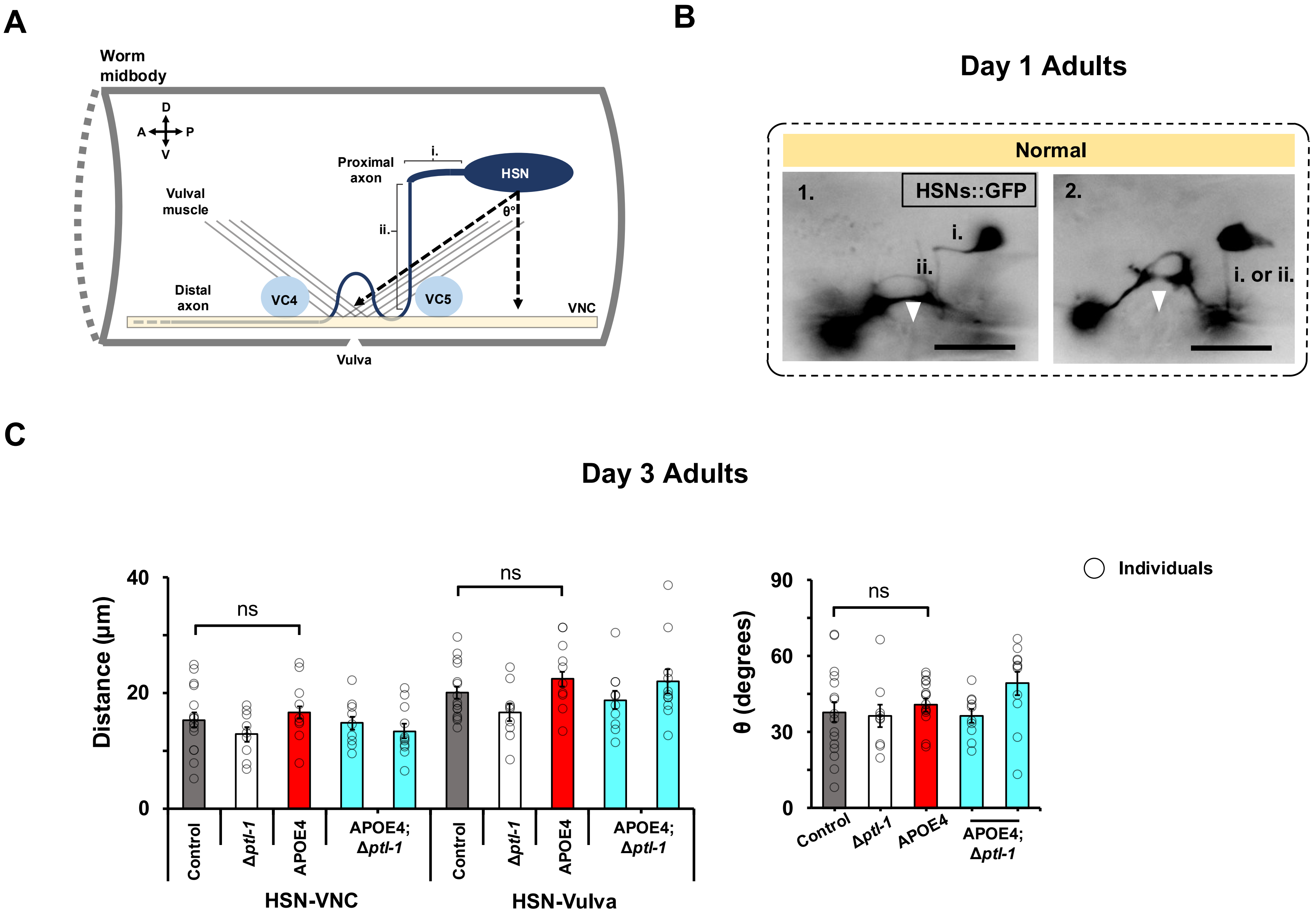
A
Worm midbody
D
A
P
V
Proximal axon
i.
HSN
Vulval muscle
ii.
θ°
Distal axon
VC4
VC5
VNC
Vulva
B
Day 1 Adults
Normal
1.
HSNs::GFP
2.
i.
ii.
i. or ii.
C
Day 3 Adults
Individuals
Distance (µm)
40
20
0
ns
ns
Control
Δptl-1
APOE4
APOE4; Δptl-1
Control
Δptl-1
APOE4
APOE4; Δptl-1
HSN-VNC
HSN-Vulva
θ (degrees)
90
60
30
0
ns
Control
Δptl-1
APOE4
APOE4; Δptl-1

### Fig. S8

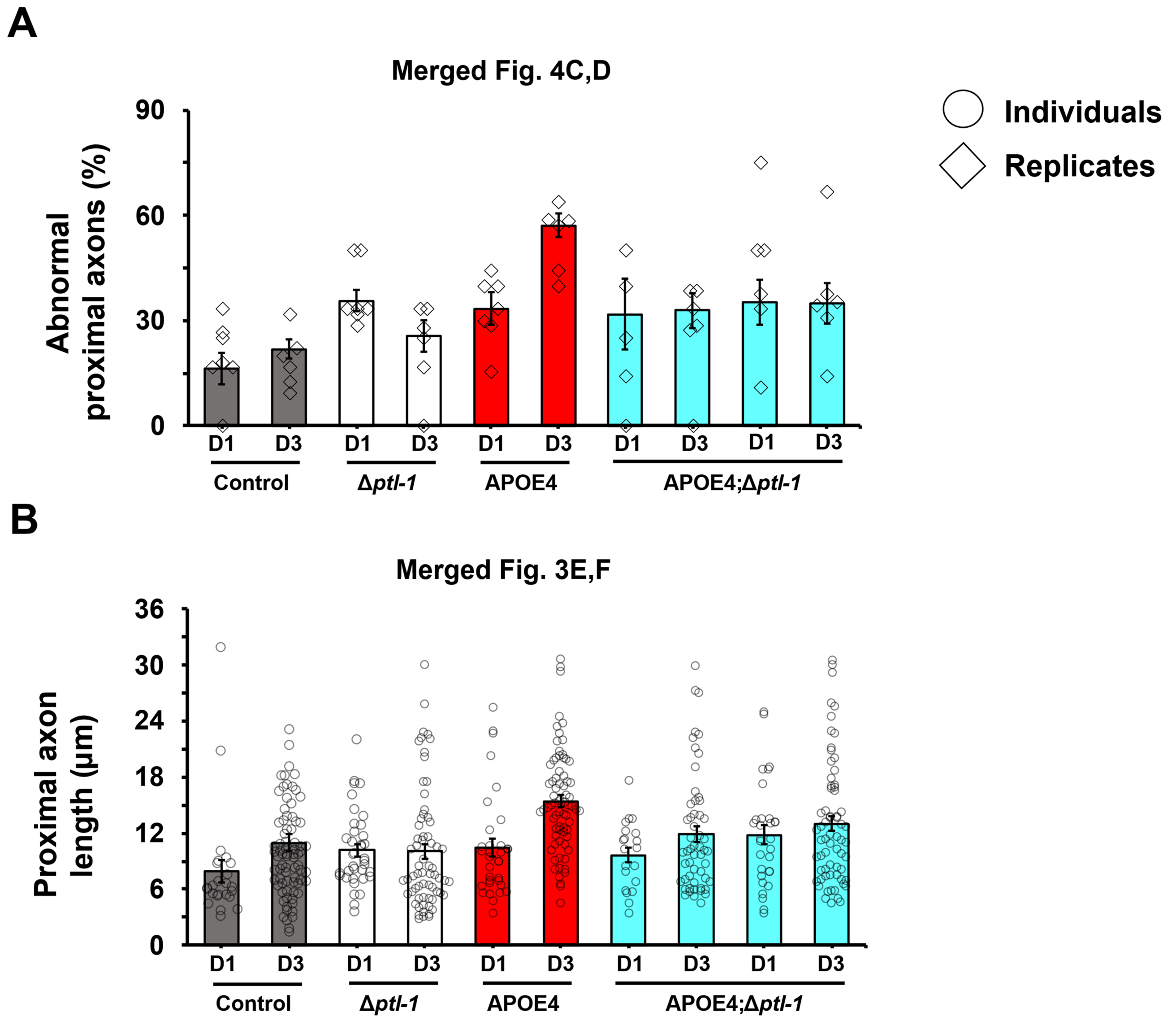
A
Merged Fig. 4C,D
Individuals
Replicates
Abnormal proximal axons (%)
90
60
30
0
D1
D3
D1
D3
D1
D3
D1
D3
D1
D3
Control
Δptl-1
APOE4
APOE4;Δptl-1
B
Merged Fig. 3E,F
Proximal axon length (µm)
36
30
24
18
12
6
0
D1
D3
D1
D3
D1
D3
D1
D3
D1
D3
Control
Δptl-1
APOE4
APOE4;Δptl-1

### Fig. S9

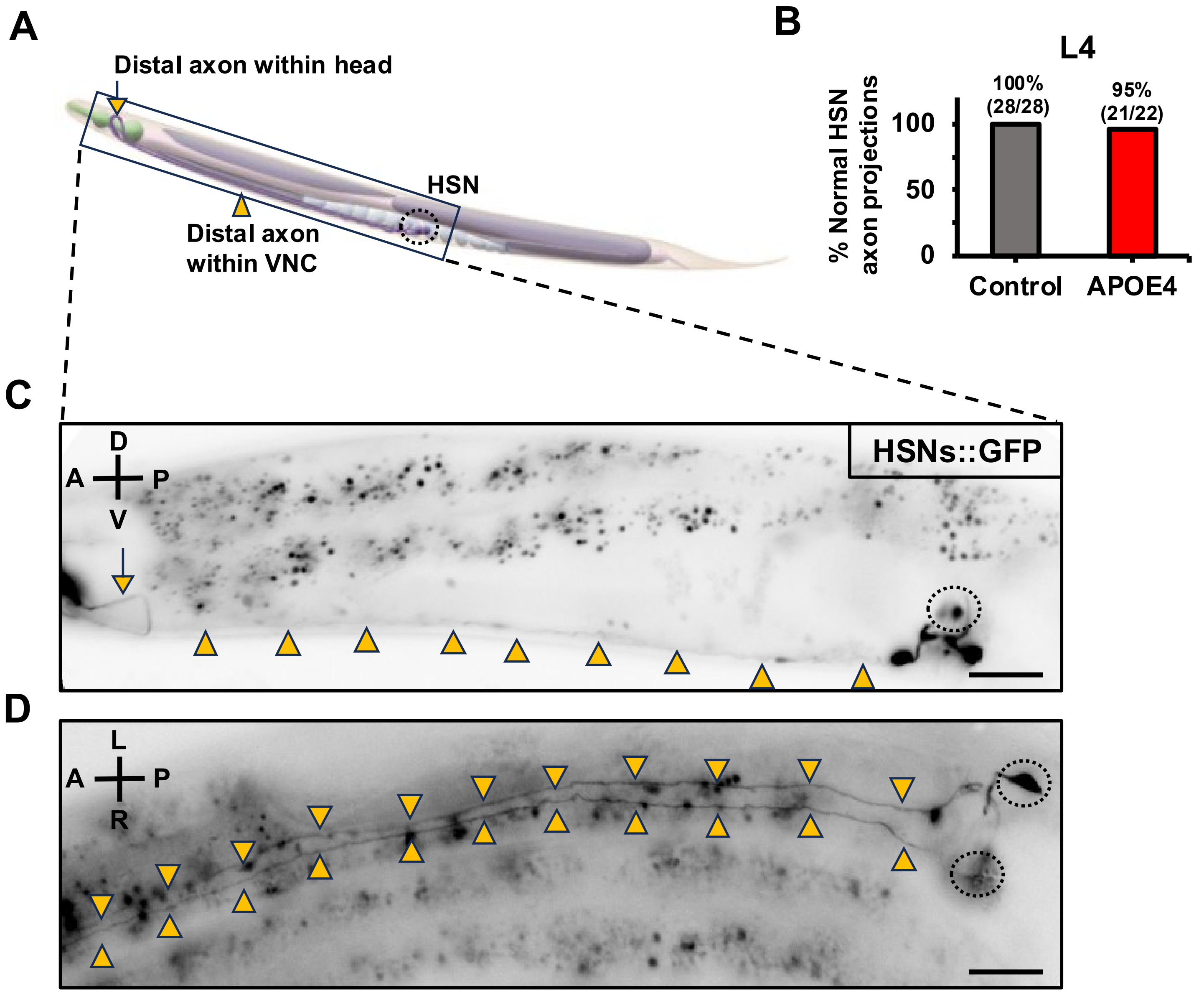
A
Distal axon within head
HSN
Distal axon
within VNC
B
L4
% Normal HSN
axon projections
100%
(28/28)
95%
(21/22)
100
50
0
Control
APOE4
C
HSNs::GFP
D
A
P
V
D
L
A
P
R

Fig. S10

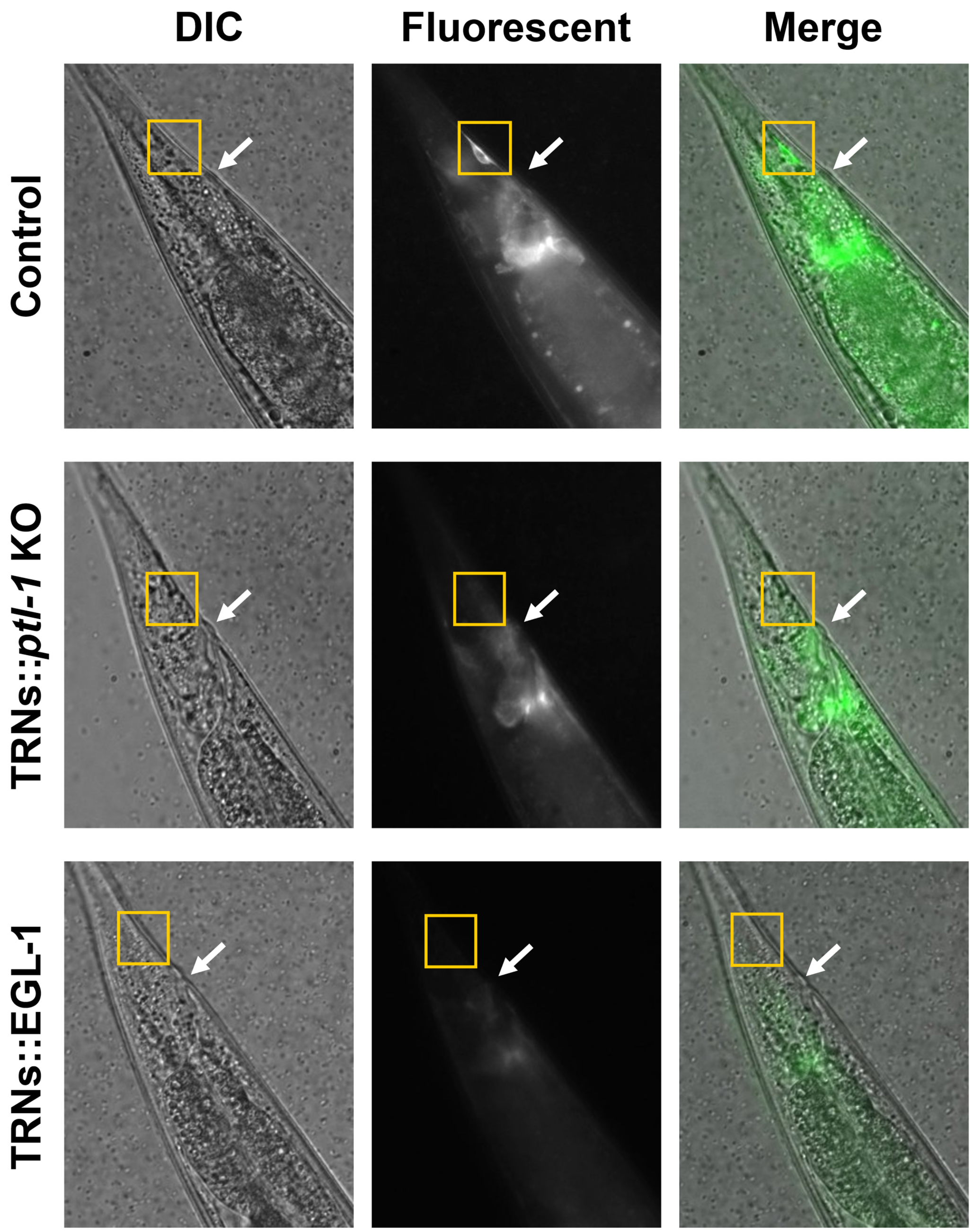

### Fig. S11

A

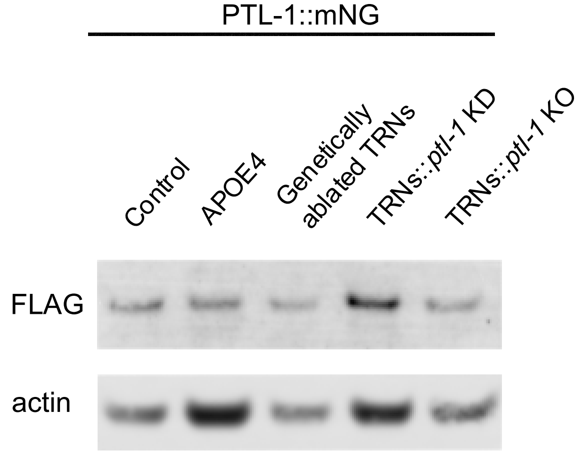
PTL-1::mNG
Control
APOE4
Genetically ablated TRNs
TRNs::ptl-1 KD
TRNs::ptl-1 KO
FLAG
actin

B

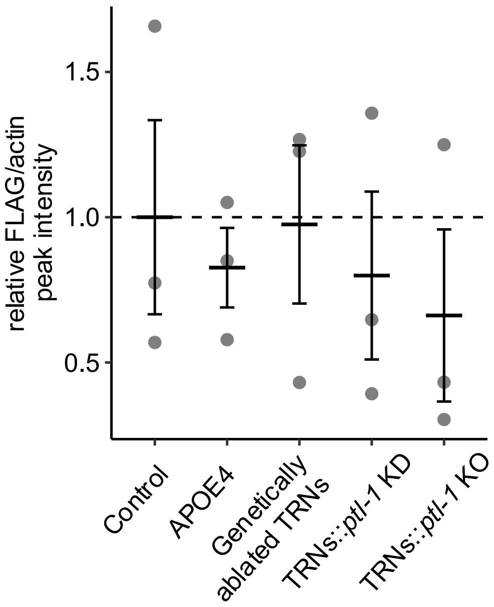
relative FLAG/actin peak intensity
1.5
1.0
0.5
Control
APOE4
Genetically ablated TRNs
TRNs::ptl-1 KD
TRNs::ptl-1 KO

### Fig. S12

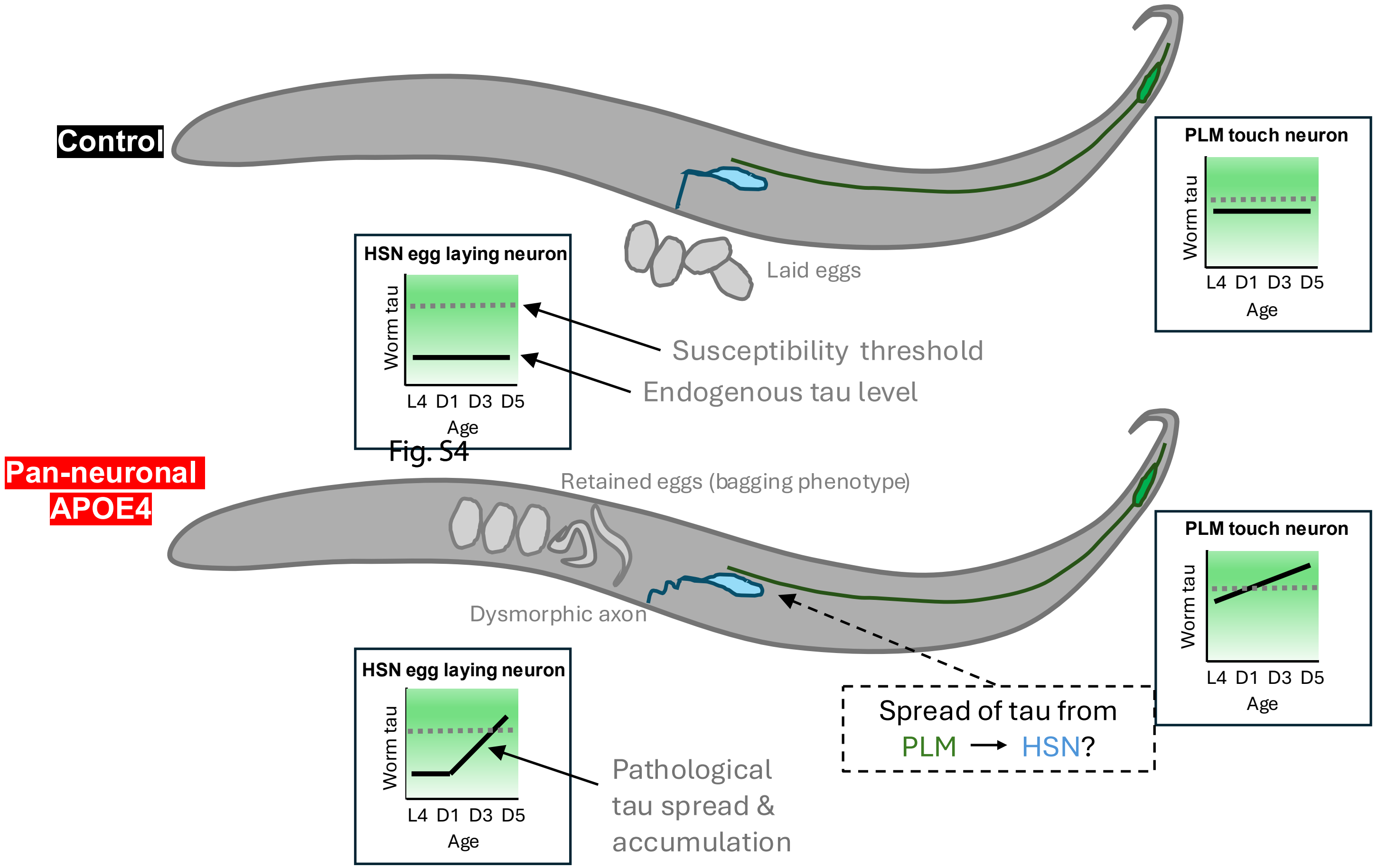
Control
PLM touch neuron
Worm tau
L4 D1 D3 D5
Age
Laid eggs
HSN egg laying neuron
Worm tau
L4 D1 D3 D5
Age
Susceptibility threshold
Endogenous tau level
Fig. S4
Pan-neuronal
APOE4
Retained eggs (bagging phenotype)
Dysmorphic axon
PLM touch neuron
Worm tau
L4 D1 D3 D5
Age
HSN egg laying neuron
Worm tau
L4 D1 D3 D5
Age
Pathological
tau spread &
accumulation
Spread of tau from
PLM → HSN?
